# Endothelial NMDA receptor involvement in retinal neurovascular damage following prenatal alcohol exposure in mouse model

**DOI:** 10.64898/2025.12.19.695085

**Authors:** Anaïs Leroy, Audrey Valentin, Camille Sautreuil, Gilles Carpentier, Camille Racine, Maryline Lecointre, François Janin, Denis Vivien, Stéphane Marret, Serge Picaud, Bruno J Gonzalez, Carole Brasse-Lagnel

## Abstract

Prenatal alcohol exposure (PAE) induces neurodevelopmental damage leading to fetal alcohol spectrum disorders (FASD) by altering both brain and ocular development. Recent data showed that PAE impairs brain cortical and retinal vasculature leading to defective positioning of interneurons. In the retina, PAE disturbs vascular development and the association of calretinin neurons with vessels. The NMDA receptor (NMDAR) is a major target of alcohol in the brain, and both ligand binding to NMDARs and the expression of NMDAR subunits are altered in FASD. Given that NMDAR is also expressed in endothelial cells and that glutamate stimulation of endothelial NMDAR (eNMDAR) regulates cortical interneuron positioning along blood vessels, we hypothesize that eNMDAR is critical for retinal vascular development and mediates PAE-induced defects.

Using an *in vivo* model of FASD and transgenic mice lacking, specifically in endothelial cells, the GluN1 subunit of the NMDAR, this study aimed to characterize the neurovascular phenotype of the developing retina. The results show that eNMDAR knockout delays the formation of the superficial vascular plexus and prevents the alterations in vascular organization and neuronal density induced by PAE, particularly cells positioned closer to the retinal vasculature, namely ganglion, amacrine, and horizontal cells. Moreover, eNMDAR deletion led to an increased number of calretinin-positive interneurons in contact with perforating vessels and prevents the decrease induced by PAE. Together, these findings demonstrate that eNMDARs are essential for normal retinal neurovascular development and mediate, at least in part, the detrimental effects of ethanol exposure in FASD.

**Significance statement:** Using a murine model of Fetal Alcohol Spectrum Disorder (FASD) and transgenic mice lacking the GluN1 subunit of the NMDA receptor specifically in endothelial cells (eNMDAR), this study demonstrates that eNMDAR plays a critical role in mediating prenatal alcohol exposure (PAE)-induced neurovascular abnormalities in the retina. Loss of eNMDAR alters the progression of the superficial vascular plexus and prevents the vascular impairments typically observed following PAE. In addition, eNMDAR deletion protects against PAE-induced neuronal damage, particularly affecting retinal ganglion cells, calbindin-positive, and calretinin-positive interneurons. Notably, this study identifies, for the first time, a role for endothelial NMDAR in regulating neurovascular interactions between retinal vessels and calretinin-positive neurons, highlighting this receptor as a key molecular mediator of ethanol-induced retinal damage.

## Introduction

Prenatal alcohol exposure (PAE) leads to fetal alcohol syndrome (FAS) characterized by abnormalities such as intrauterine growth retardation, distinct craniofacial dysmorphism, and central nervous system (CNS) anomalies, resulting in cognitive and behavioral deficits (Burd et al., 2003). FAS is the leading cause of acquired, non-genetic intellectual disability in industrialized countries (Burd et al., 2003, Popova et al., 2017). Many individuals with fetal alcohol spectrum disorder (FASD) remain undiagnosed until neurodevelopmental symptoms manifest in childhood (Chasnoff et al., 2015). Early diagnosis, ideally prior to the onset of neurodevelopmental disorders, is crucial for implementing clinical care during peak neuroplasticity (Lerna et al., 2014). A better understanding of the pathophysiological mechanisms underlying FASD could help identify new diagnostic markers and potentially open up novel therapeutic avenues.

Clinical and preclinical studies have shown that neurobehavioral disabilities observed in FASD are associated with impaired development in various brain structures, differentiation and positioning of different cell types, such as GABAergic interneurons (Léger et al., 2020a) or oligodendrocytes (Brosolo et al., 2022; Treit et al., 2013; Inkelis et al., 2020). PAE also affects ocular development, with visual problems being more prevalent than in the general population (Strömland et al., 1987). Recent studies indicate that PAE disrupts retinal vascular development and the positioning of calretinin-positive cells (Dumanoir et al., 2023). However, molecular mechanisms underlying PAE’s effects on neurovascular development in retina remain poorly understood.

The NMDA receptor (NMDAR) is a key target of ethanol in the CNS (Hoffman et al., 1990, Chandrasekar, 2013) and plays a role in ethanol-related phenotypes including dependence, withdrawal and relapse (Holmes et al., 2013). Ethanol modulates NMDAR by altering ion channel gating, reducing channel opening frequency and mean open time (Lima-Landman and Albuquerque, 1989). Indeed, studies by Ren et al. (2007, 2012) and Zhao et al. (2016) have shown that ethanol interacts with GluN1 and GluN2 subunits. Altogether, these data suggest that NMDAR contributes to ethanol impacts in FASD. However, due to the broad expression pattern of NMDAR in different cell types including neurons (Paoletti et al., 2013), astrocytes (Kosenkov et al., 2024), oligodendrocytes (Salter and Fern, 2005), and microglial cells (Raghunatha et al., 2020), deciphering the pathophysiological contribution of the endothelial NMDAR (eNMDAR) in FASD remains challenging.

Previous studies indicated that cortical endothelial cells express NMDAR during perinatal development (Legros et al., 2009; Léger et al., 2020b). In particular, it has been shown that glutamate, by stimulating MMP9 activity through an endothelial NMDAR-dependent mechanism, stimulates the migration of GABA interneurons along vessels (Léger et al., 2020b). Interestingly, in a mouse model of FASD, cortical vascular dysfunction was associated with a strong overexpression of the GluN1 subunit, increased MMP-9 activity, and mispositioning of Gad67-GFP interneurons, which normally populate superficial cortical layers (Léger et al., 2020a). Taken together, these studies suggest a role for the endothelial NMDAR (eNMDAR) in the recently reported effects of PAE on retinal vascular development and interneuron-vessel interactions (Dumanoir et al., 2023). Consistent with this hypothesis, intravitreal injection of NMDA in rat neonates led to neuronal cell loss and retinal capillary degeneration (Asano et al., 2019). Therefore, exploring the role of eNMDAR in retinal neurovascular development could help decipher mechanisms involved in the deleterious effects of ethanol on CNS angiogenesis and provide new research avenues for the early diagnosis of FASD.

This study aims to define the role of eNMDAR in neurovascular development of retina and its contribution to the effects of PAE. Using a mouse model of FASD in both wild-type and endothelial-specific Grin1 knockout mice, we characterized the impact of eNMDAR loss on (i) the development of superficial and deep vascular plexuses, (ii) neuronal density across retinal layers and (iii) the vessel association of interneurons.

## Materials and Methods

### Animals

All animals used were from the same genetic background (C57BL6J). C57BL/6J mice were sourced from Janvier (Le Genest-Saint-Isle, France). Floxed Grin1 mice (Grin1^fl/fl^; B6.129S4-Grin1tm2Stl/J; #005246) and VE-Cadherin-CRE mice (VE-Cad^CRE^; B6.FVB-Tg(Cdh5-cre)7Mlia/J; #006137) were obtained from The Jackson Laboratory (Bar Harbor, Maine, USA). VE-Cad^CREΔGrin1^ mice were generated by our group by crossing Grin1^fl/fl^, in which exon 11-21 region of the Grin1 gene was flanked by loxP sites, with VE-Cad^CRE^. This study was conducted on pups obtained by crossing homozygous Grin1^fl/fl^ female with homozygous VE-Cad^CREΔGrin1^ male. This breeding strategy allows for Cre-mediated germline excision of exons 11 to 21 of the endothelial *Grin1* gene, resulting in the deletion of both alleles in the offspring VE-Cad^CREΔGrin1^. Animals were housed in temperature-controlled rooms (21 ± 1°C) with a 12-hour light/dark cycle and had free access to food and water. All animal procedures were performed in accordance with the European Communities Council Directives (2010/63/EU) and French National legislation (ethical approval no. 01680.02) under the supervision of authorized personnel (CBL authorization no. 37567 from the French Ministry of Agriculture and Fisheries).

### Genotyping by PCR

Mutant mice were genotyped by PCR using the following primers: for Grin1, 5′-AAACAGGGCTCAGTGGGTAA and 3′-GTGCTGGGATCCACATTCAT; for the Cre transgene, 5′-GCGGTCTGGCAGTAAAAACTATC and 3′-GTGAAACAGCATTGCTGTCACTT. Genotypes were determined based on distinct band shifts corresponding to mutation size, visualized on a 3.5% agarose gel (Léger et al., 2020b).

### Chemicals

Bovine serum albumin (BSA), ethanol, glycerol, Hoechst, phosphate-buffered saline (PBS), sucrose, NaCl (0,9%) and Triton X-100 were obtained from Sigma-Aldrich (Saint Quentin Fallavier, France). Paraformaldehyde (PFA) was obtained from Labonord (Templemars, France). Vetflurane was purchased by Baxter (Maurepas, France). MK801 was obtained from Tocris (R&D, Lille, France). Epredia™ Neg-50™ Frozen Section Medium (Neg-50™) and PermaFluor™ were obtained from Thermo Fisher Scientific France (Illkirch-Graffenstaden, France). The complete mouse endothelial cell culture medium (M1168) was obtained from Cell Biologics (Chicago, IL, USA). Matrigel® Matrix was from Corning (New York, USA) and μ-Slide angiogenesis dish from Ibidi (Ibidi, Munich, Germany). References, providers and dilutions of the primary antibodies against Calbindin, Calretinin, CD31, GluN1, Opsin B, Opsin R/G, RBPMS and Rhodopsin are listed in Table 1. The secondary antibodies used for immunohistochemistry are also listed in this Table 1.

**Table 1:** Antibodies. Origin and characteristics of immunohistochemical antibodies.

| Primary antibodies<br>(dilution) | Provider/Reference | Secondary Antibody<br>(dilution) | Provider/Reference |
| --- | --- | --- | --- |
| Rat anti-CD31<br>(1/200) | BD Pharmingen<br>/550274 | Donkey Alexa Fluor® 594 anti-rat IgG (1/400) | Molecular<br>Probes/A21209 |
| Rabbit anti-Calbindin<br>(1/200) | Abcam/ab108404 | Donkey Alexa Fluor® 488 anti-<br>rabbit IgG (1/400) | Invitrogen/A-21206 |
| Rabbit anti-Calretinin<br>(1/500) | Thermofisher<br>Scientific/180211 |  |  |
| Rabbit anti-RBPMS<br>(1/500) | Merck<br>Millipore/ABN1362 |  |  |
| Rabbit anti-opsin B<br>(1/500) | Merck<br>Millipore/AB5407 |  |  |
| Rabbit anti-opsin RG<br>(1/500) | Merck<br>Millipore/AB5405 |  |  |
| Rabbit anti-rhodopsin<br>(1/500) | Abcam/ab112576 |  |  |
| Rabbit anti-GluN1<br>(1/200) | Abcam/ab17345 |  |  |

### In vivo treatment of pregnant mice and newborns

Females were placed at 4pm with VE-Cad^CREΔGrin1^ males (E0) and separated the following day at 8am and gestation was verified by follow-up weighing. Pregnant mice received daily subcutaneous injections of 0.9% NaCl or ethanol (3 g/kg), diluted 50% (v/v) in 0.9% NaCl, from gestational day 14 (GD14) until parturition (P0). The injection volume was adjusted daily according to the pregnant mouse’s weight. Experiments on neonates were conducted at P5 and P15, two developmental stages shown to be impacted by PAE regarding retina neurovascular development (Dumanoir et al., 2023). Pups were randomly selected to maintain uniform litter sizes. Both male and female pups were used in the experiments and a corresponding littermate colony (colonies maintained and treated with NaCl in parallel) was used as a control for each genotype.

### Whole-mount retina immunohistochemistry

Mice from control and ethanol-exposed groups were euthanized at P5 and P15 using Vetflurane (5%), followed by decapitation. Eyes were collected and fixed for 30 minutes in 4% paraformaldehyde (PFA) at room temperature under agitation. Retinas were then isolated, blocked and immunostained with a rat anti-CD31 antibody (1:200) overnight at 4°C in a solution containing 0.2% BSA and 0.5% Triton X-100 in PBS. Secondary staining was performed for 1.5 hours at room temperature with Alexa Fluor® 594 donkey anti-rat IgG (1:400 in 0.2% BSA and 0.5% Triton X-100 in PBS) under agitation. Retinas were mounted with PermaFluor™ and imaged using a THUNDER Imaging System (Leica, Reuil-Malmaison, France). Morphometric analysis of the whole-mount retinas was performed using ImageJ (National Institutes of Health, Bethesda, MD) software with the *Angiogenesis Analyzer* plugin (Carpentier et al., 2012 and 2020).

### Cross sections of retinas and immunostaining

Eyes fixed in 4% PFA were incubated in PBS/sucrose gradients (10%, 20%, 30%), dissected, embedded in Neg-50™, and frozen. Serial 20 µm sections were prepared using a cryostat (Leica CM1950) and used for immunohistochemical labeling. Sections were blocked and incubated with primary antibodies for vessels and specialized neuronal cell types, followed by secondary antibodies using the same procedure as previously described for whole-mount retinas (Table 1; Dumanoir et al., 2023). Cell nuclei were stained with Hoechst 33258 (1:5000) for 5 minutes before mounting in a PBS/Glycerol (50/50 v/v) solution. Imaging and quantification were performed using THUNDER Imaging Systems or an SP8 confocal microscope with *LAS AF Lite software* (Leica). Quantified parameters included the number of positive cells per 100 µm, the number of perforating vessels per 100 µm and the number of calretinin-positive cells associated with vessels.

### Endothelial cell culture

Primary brain microvascular endothelial cells (BMVECs) from C57BL/6J mouse pups (P2) were obtained from Cell Biologics (C57-6023). These cells were cultured at 37°C in a humidified environment with 5% CO₂. They were maintained in a complete mouse endothelial cell culture medium (M1168), fed every 2 days, and grown on 60-mm dishes. Upon reaching confluence, the cells were incubated overnight in serum-free medium and treated with ethanol (50 mM), and/or MK-801 (dizocilpine maleate, 20 µM).

### Tube formation assay in matrigel

The tube formation assay was conducted according to the manufacturer’s instructions. On the day of the assay, 10 μl of Matrigel® Matrix was added to an Ibidi μ-Slide angiogenesis dish and incubated at 37°C with 5% CO₂ for 0.5 hours. During this incubation, BMVECs treated overnight with or without 20 µM MK-801 in serum-free medium were harvested and counted. A cell suspension (30 μl) containing 5000 cells was added to each well with Matrigel® Matrix. The slides were then incubated at 37°C for 5 hours with or without 50 mM ethanol and/or 20 µM MK-801, and tube formation was visualized and captured using a Zeiss 200M phase-contrast microscope at ×20 magnification. Tube formation was quantified by measuring the number of meshes using ImageJ software with the *Angiogenesis Analyzer* plugin. Each condition was tested in triplicate. The control condition was normalized to a value of 1, and experimental values were expressed as ratios relative to the control.

### Statistical analysis

Statistical analyses were performed using GraphPad Prism version 9 for Windows (GraphPad Software). Data from male and female mice were pooled for these analyses. As the sample size per group was small (n < 10), normality tests lacked sufficient power. Non-parametric methods, which do not require normally distributed data, were therefore used for statistical analysis. So, a non-parametric one-way ANOVA (Kruskal–Wallis test) was used to assess differences between groups. When the Kruskal–Wallis test indicated a significant overall effect (p < 0.05), pairwise comparisons were subsequently performed using Mann–Whitney U tests. These post hoc analyses aimed to identify significant differences among the mouse strains within the control groups, and to specifically compare the PAE group to its strain-matched control group. A ratio paired t-test was used to analyze data obtained from cultured cells. Results from mouse experiments are presented as box plots showing the median, minimum, and maximum values, while data from cultured cells are presented as bar graphs representing the mean ± standard error of the mean (SEM). Statistical significance was defined as p ≤ 0.05. All statistical results are summarized in Table S1.

## Results

### Endothelial NMDAR knockout delays retinal vascular development and prevents several PAE-induced impairments during early angiogenesis

To determine whether eNMDAR is involved in the *in vivo* effects of ethanol on retinal vascular development, experiments were conducted on VE-Cad^CREΔGrin1^ transgenic mice exposed *in utero* to alcohol. GluN1 is an essential subunit required for the assembly and function of NMDA receptors. Its deletion in the VE-Cad^CREΔGrin1^ mice results in the expression of non-functional NMDA receptors in endothelial cells. These experiments were conducted in offspring obtained by crossing VE-Cad^CREΔGrin1^ homozygous males with Grin1^fl/fl^ homozygous females. As Cre-recombinase expression, controlled by the VE-Cadherin promoter, occurs as early as E7.5 (Alva et al., 2006), the Grin1 gene is specifically invalidated in endothelial cells of VE-Cad^CREΔGrin1^ transgenic pups during the period of *in utero* alcohol exposure in the FASD mouse model-i.e. during the third week of gestation-as well as at the developmental stages analyzed (P5 and P15). As shown in Figure 1, GluN1 is expressed in both nervous cells and blood vessels within the retinal tissue of wild type mice (Fig. 1A (a-k)) and its expression is specifically lost in the vasculature of VE-Cad^CREΔGrin1^ mice (Fig. 1B (a-k)). Indeed, as revealed by IMARIS analysis, a significant loss of CD31/GluN1 colocalization was observed in the retinal vessels of VE-Cad^CREΔGrin1^ mice (Fig. 1A (l-m) *vs* 1B (l-m)).

**Figure 1.**
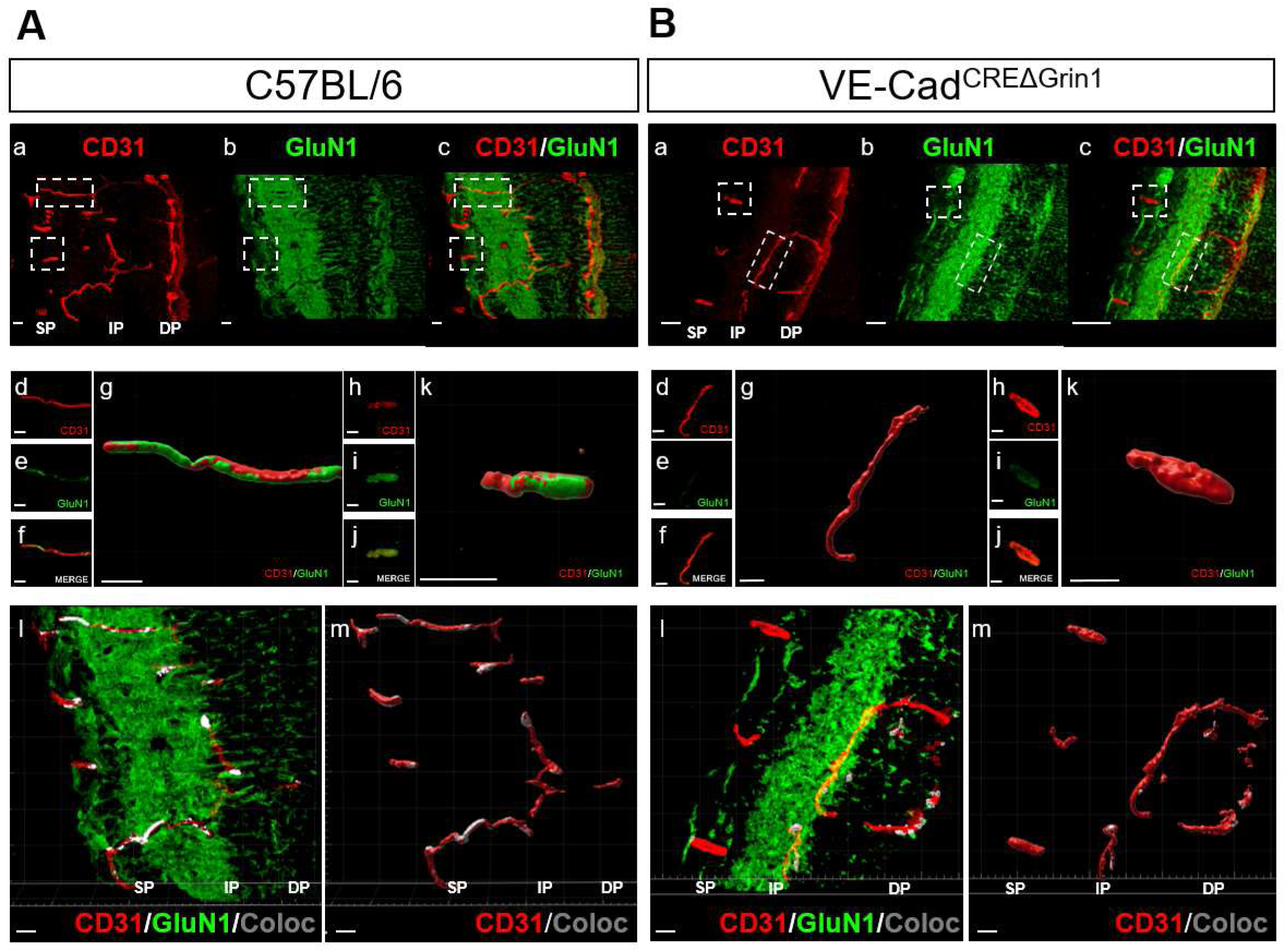
Endothelial GluN1 expression loss in transgenic mice VE-Cad^CREΔGrin1^. **(A-B)** Sagittal sections of C57BL/6J (A) and VE-Cad^CREΔGrin1^ (B) retinas at P15 were immunolabelled against CD31 (revealed with Alexa 594 in red) and GluN1 (revealed with Alexa 488 in green) and observed by using confocal microscopy. (a-c) CD31-positive microvessels (a) and GluN1 immunoreactivity (b) were visualized in the superficial (SP), intermediate (IP), and deep (DP) vascular plexuses. The overlay is shown in c. (d-k) Delineation by using IMARIS thresholding of CD31- (d, h) and GluN1- (e, i) immunostaining in two microvessels from the regions of interest outlined in a-c. The overlay of both signals is visualized in f and j. 3D modelling is illustrated in g and k. (l-m) Colocalization analysis performed using the IMARIS “Coloc” module of CD31 and GluN1 immunostainings. The colocalized signal is highlighted in gray in confocal images (l) and in the 3D modelling (m). For all these analyses, the threshold and modelling parameters were firstly determined in C57BL/6J-related images and then applied to VE-Cad^CREΔGrin1^-related images. Scale bars: 10 µm

We next examined how eNMDAR deletion affects both basal and alcohol-induced retinal angiogenesis in P5 mice, a developmental stage characterized by active vessel growth. At P5, the superficial vascular plexus is still developing, whole-mount retina immunofluorescent staining for CD31 showed delayed progression of the superficial vascular network in VE-Cad^CREΔGrin1^ mice compared to C57BL/6J, Grin1^fl/fl^ and VE-Cad^CRE^ control mice (Fig. 2A (a-d) and B (hatched white box *vs* solid white boxes)). Quantification showed a significant increase in distance to be covered in VE-Cad^CREΔGrin1^ mice: +34% compared to C57BL/6J (p=0.0057, n=10-9, Mann Whitney U test), +33% compared to Grin1^fl/fl^ (p=0.0030 n=10-9, Mann Whitney U test) and +21% compared to VE-Cad^CRE^ (p=0.0159, n=11-9, Mann Whitney U test). Similarly, quantification of vascular density revealed a significant increase in VE-Cad^CREΔGrin1^ mice compared to the Grin1^fl/fl^ and VE-Cad^CRE^ control groups (Fig. 2C (a-d) and 2D (hatched white box *vs* solid white boxes); +16% compared to Grin1^fl/fl^, p=0.0016, n=9-8; +15% compared to VE-Cad^CRE^, p=0.0155, n=10-8, Mann Whitney U test). However, no significant differences were observed in the number of meshes or in segment length between the control groups (C57BL/6J, Grin1^fl/fl^, and VE-Cad^CRE^) and VE-Cad^CREΔGrin1^ transgenic mice (Fig. 2E and 2F (hatched white box *vs* solid white boxes)). Altogether, these data suggest that at P5, eNMDAR knockout alters the progression of the superficial vascular plexus without affecting the overall vascular morphology.

**Figure 2.**
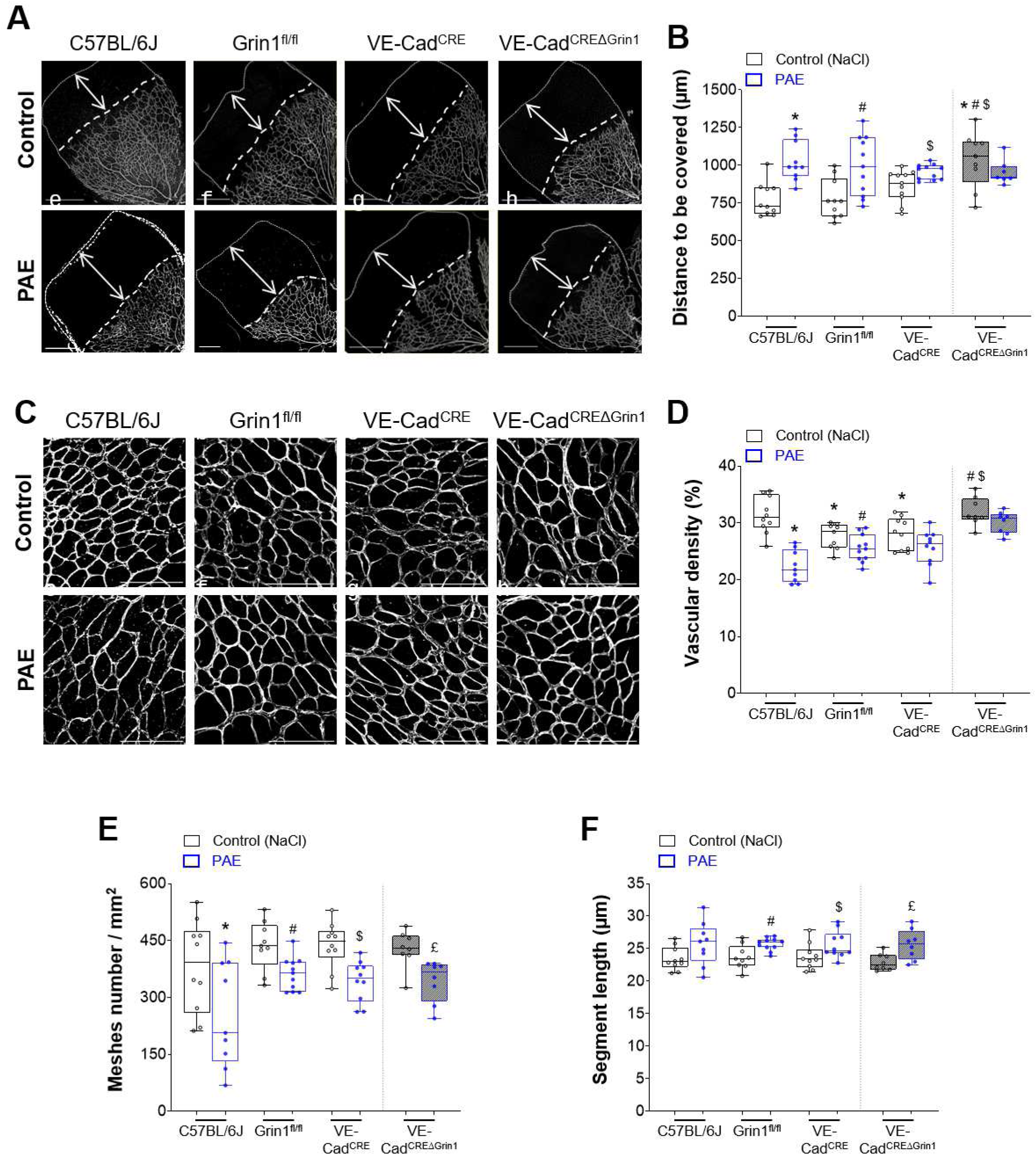
Effect of eNMDAR knockout on retinal vascular development and PAE-induced damage at postnatal day 5. **(A)** Representative images of the superficial vascular plexus in whole-mount retinas at P5, following CD31 immunostaining. Retinas were collected from control (NaCl; a–d) and PAE-exposed (e–h) neonates of full eNMDAR mice (i.e. C57BL/6J, Grin^fl/fl^ and VE-Cad^CRE^) and eNMDAR null mice (VE-Cad^CREΔGrin1^). Double-headed arrows indicate the remaining distance between the vascular front and the retinal periphery. Scale bars: 500 µm. (**B**) Quantification of the distance to be covered by the vascular plexus to reach the retinal edge in control (NaCl; white boxes) and PAE (blue boxes) conditions across all genotypes. Kruskal–Wallis test, H(7) = 28.74, p = 0.0002. (**C**) Higher magnification images showing the superficial vascular network in control (NaCl; a–d) and PAE (e–h) conditions for each genotype. Scale bars: 200 µm. (**D–F**) Quantification of vascular density (D, Kruskal–Wallis test, H(7) = 44.79, p <0.0001), mesh number (E, Kruskal–Wallis test, H(7) = 24.45, p=0.0021) and segment length (F, Kruskal–Wallis test, H(7) = 19.51, p =0.0067) at P5 in control (NaCl; white boxes) and PAE (blue boxes) retinas from C57BL/6J, Grin1^fl/fl^, VE-Cad^CRE^, and VE-Cad^CREΔGrin1^ mice. Statistical significance: *p ≤ 0.05 *vs* C57BL/6J control (NaCl); #p ≤ 0.05 *vs* Grin1^fl/fl^ control (NaCl); $p ≤ 0.05 *vs* VE-Cad^CRE^ control (NaCl); £p ≤ 0.05 *vs* VE-Cad^CREΔGrin1^ control (NaCl); Mann–Whitney test, n = 8–11 per group. *PAE: Prenatal Alcohol Exposure*.

Concerning the impact of PAE on retinal vascular development, PAE impaired or delayed vascular outgrowth across all control groups (C57BL/6J, Grin1^fl/fl^, and VE-Cad^CRE^; Fig. 2A,C (a–c; e–g) and 2B,D (blue boxes)), as evidenced by an increase in the remaining distance to be covered (Fig. 2B; +33% for C57BL/6J, p=0.0007, n=10-10; +28% for Grin1^fl/fl^, p=0.0079, n=10-11; and +12% for VE-Cad^CRE^, p=0.00336, n=11-11, Mann Whitney U test) and a reduction in vascular density (Fig2D; -29% for C57BL/6J, p<0.0001, n=10-9; -8% for Grin1^fl/fl^, p=0.0465; n=9-11; and -8% for VE-Cad^CRE^, p=0.1655, n=10-10, Mann Whitney U test). Interestingly, these alterations were not observed in VE-Cad^CREΔGrin1^ mice (Fig. 2A,C (d, h) and 2B,D (hatched blue box *vs* hatched white box; B: p=0.1738, n=9-7; D: p=0.2786, n=8-8, Mann Whitney U test). Regarding vascular morphology, PAE led to a reduction in the number of meshes (Fig. 2E; -33% for C57BL/6J, p=0.0435, n=10-9; -17% for Grin1^fl/fl^, p=0.0125, n=9-11; -19% for VE-Cad^CRE^, p=0.0029, n=10-10, Mann Whitney U test) and an increase in segment length (Fig. 2F; +10% for C57BL/6J, p=0.1333, n=10-9; +9% for Grin1^fl/fl^, p=0.0097, n=9-11; and +6% for VE-Cad^CRE^, p=0.0355, n=10-10, Mann Whitney U test) across all mouse strains, including VE-Cad^CREΔGrin1^ (Fig. 2E,F (hatched blue box *vs* hatched white box); mesh number: -20%, p=0.0047, n=8-8; segments length: +12%, p=0.0104, n=8-8, Mann Whitney U test). Complementary *in vitro* experiments using Matrigel, a well-established model that recapitulates the initial steps of angiogenesis, revealed that the ethanol-induced reduction in blood vessel formation can be prevented by NMDA receptor inhibition with MK-801 (Fig. 3). Indeed, the decrease in mesh number induced by ethanol incubation is not observed in the presence of MK801 (Fig. 3B; without MK801: -33% ± 5%, p=0.0018, n=8; with MK801: -17% ± 18%, p=0.1069, n=8, ratio paired *t* test).

**Figure 3.**
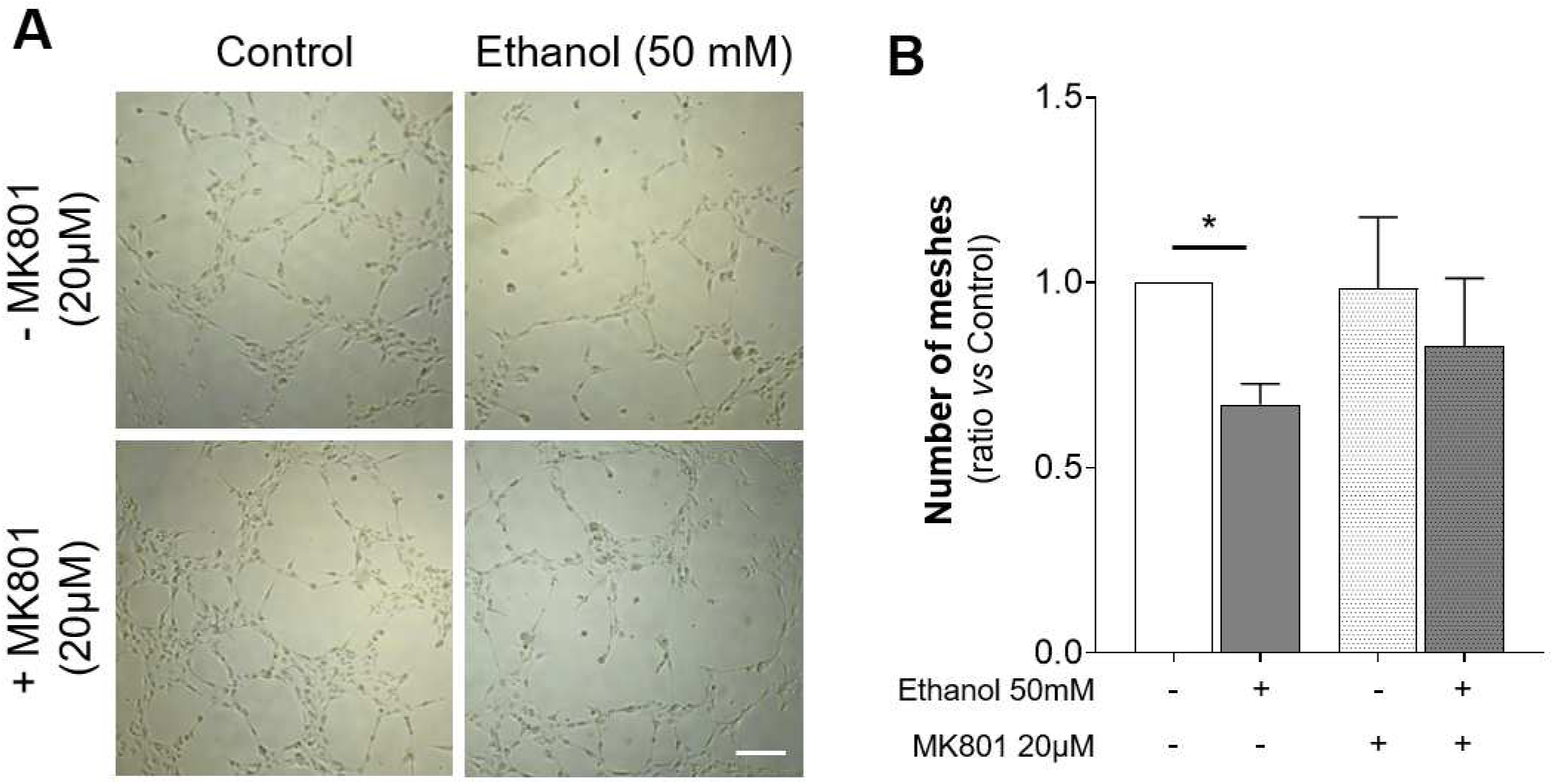
Effect of ethanol and MK801 on pseudocapillary-tube formation in a Matrigel® assay. Experiments were conducted using brain microvascular endothelial cells (BMVECs) placed on Matrigel®. Cells were treated overnight with or without NMDAR inhibitor -MK801 (20 µM)- followed by a 5-hour incubation on Matrigel® with or without ethanol 50 mM. (**A)** Representative images of pseudocapillary-tube formation under each experimental condition, captured using a Zeiss 200M wide-field microscope at 20x magnification. Scale bar : 75 µm (**B)** Quantification of mesh number in pseudocapillary-tube formation assay using ImageJ with the *Angiogenesis Analyser* plugin. Data are presented as mean ratios relative to control conditions ± SEM, from three replicates across eight independent experiments. Statistical comparisons: *p ≤ 0.05 compared with control condition, Ratio paired *t* test, n=8, control *vs* Ethanol: p=0.0124; control *vs* MK801: p=0.6821; MK801 *vs* MK801+Ethanol: p=0.1452. *MK801: dizocilpine*.

Altogether, these results demonstrate that PAE disrupts both the progression and morphology of the superficial vascular network and that the loss of eNMDAR prevents the ethanol-induced reduction in vascular network progression and density without affecting its morphological alterations. This suggests that eNMDAR is involved in the molecular mechanisms of early stages of angiogenesis, playing a role in network progression rather than branching.

### Endothelial NMDA receptor knockout prevents several PAE-induced impairments on retinal vasculature at P15

The effects of eNMDAR knockout and PAE on retinal vascular development were analyzed at P15 in the superficial and deep vascular plexuses across both central and peripheral regions of the retina. Under control conditions, morphometric analysis of the superficial vascular plexus revealed strain specificities on the morphology of either the superficial or deep plexus (Fig. S1). Consequently, to facilitate interpretation regarding the effects of PAE, the results were then expressed as a percentage relative to the mean of the strain-matched control group and figure 4 presents data from C57BL/6J and VE-Cad^CREΔGrin1^ transgenic mice. Results from the additional control groups (i.e., Grin1^fl/fl^ and VE-Cad^CRE^) are fully presented in the supplementary figures S2 and S3 (Fig.S2-S3).

**Figure 4.**
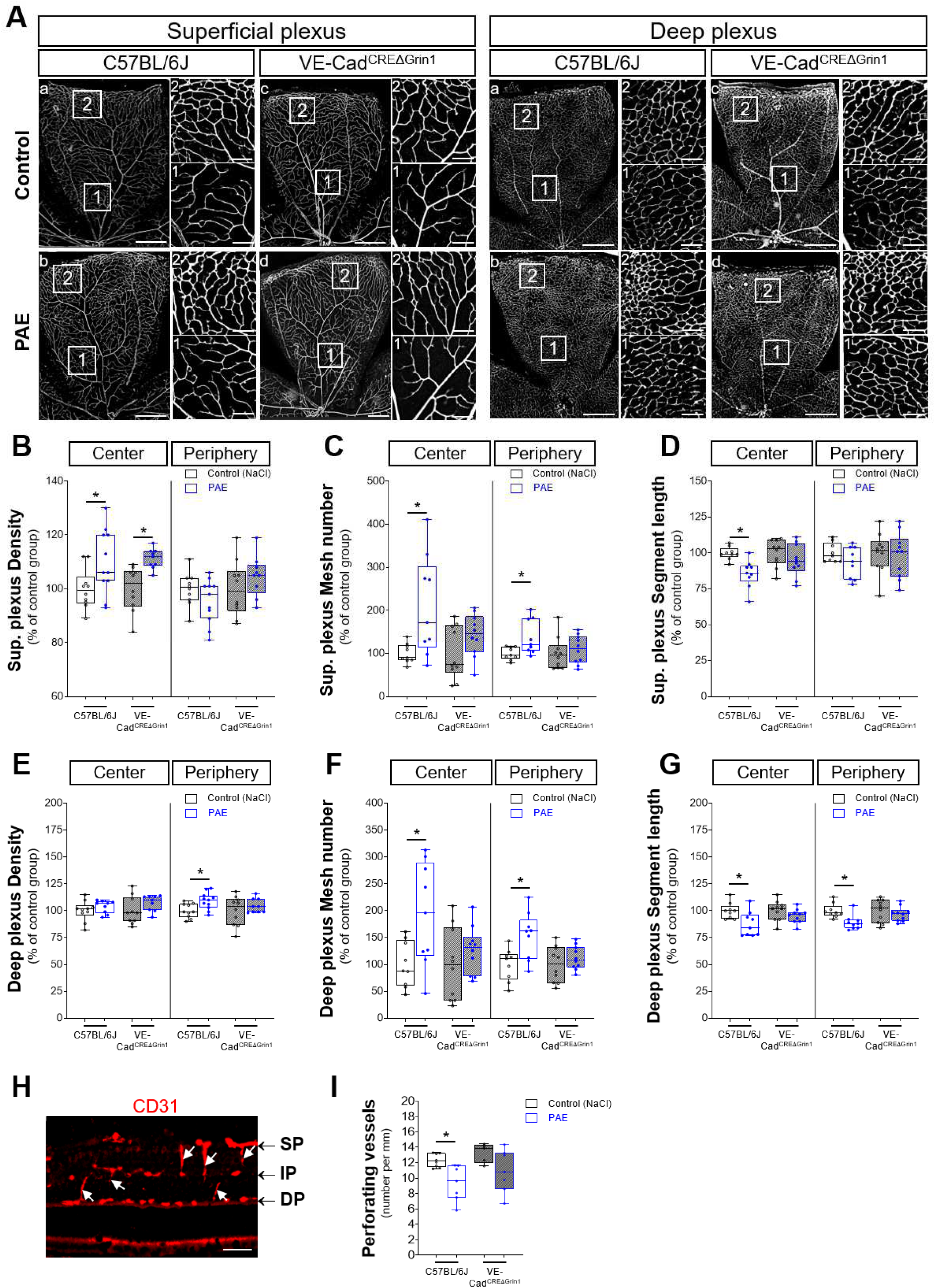
Effect of eNMDAR knockout on PAE-induced alterations in the superficial and deep retinal vascular networks at postnatal day 15. (**A**) Representative images of the superficial and deep vascular plexuses in whole-mount retinas at P15, following CD31 immunostaining. Retinas were collected from control (NaCl; a,c) and PAE-treated (b,d) C57BL/6J and VE-Cad^CREΔGrin1^ mice. Scale bars: 500 µm. Insets show high-magnification views of the superficial vascular network in the central (1) and peripheral (2) retina. Scale bars: 100 µm. (**B–G**) Quantification of vascular density (B,E), mesh number (C,F) and segment length (D,G) in central and peripheral retina regions of the superficial (B-D) and deep (E-G) plexuses at P15. Results are expressed as a percentage relative to the mean of the strain-matched control group. Statistical significance: *p ≤ 0.05 *vs* strain-matched control group (NaCl); Mann–Whitney test, n = 7-11 per group. *PAE: Prenatal Alcohol Exposure.* (**H**) Representative image of CD31 immunostaining in a cross section of P15 retina. Arrows indicate perforating microvessels. Scale bar: 100 µm. (**I)** Quantification of perforating microvessels per mm in the central retina of C57BL/6J (solid boxes) and VE-Cad^CREΔGrin1^ (hatched boxes) mice under control (NaCl, white boxes) and PAE (blue boxes) conditions. Kruskal–Wallis test, H(3) = 41.23, p = 0.0204. Statistical symbols: *p≤0.05 compared with C57BL/6J control (NaCl) group; Mann-Whitney test, n=6-8 pups per group. *DP: Deep vascular Plexus; IP: Intermediate vascular Plexus; PAE: Prenatal Alcohol Exposure; SP: Superficial vascular Plexus*.

As shown in Figure 4, in C57BL/6J mice, particularly in the central retina, PAE significantly altered vessel density, mesh number, and segment length of the superficial plexus (Fig. 4B-D, blue box *vs* solid white box). Several of these PAE-induced effects were abrogated in VE-Cad^CREΔGrin1^ mice (Fig. 4B-D, hatched blue box *vs* hatched white box). Indeed, whereas eNMDAR deletion had no significant impact on the effect of PAE on vascular density in the central retina (Fig. 4B; +10% for C57BL/6J, p=0.0430, n=10-11 and +11% for VE-Cad^CREΔGrin1^, p=0.0015, n=10-9, Mann Whitney U test), it prevented the PAE-induced effects on mesh number (Fig. 4C; +110% for C57BL/6J, p=0.0188, n=9-9 and +42% for VE-Cad^CREΔGrin1^, p=0.1431, n=10-10 for VE-Cad^CREΔGrin1^, Mann Whitney U test) and segment length (Fig. 4D; -15% for C57BL/6J, p=0.0012, n=9-9 and -5% for VE-Cad^CREΔGrin1^, p=0.3527, n=10-10 for PAE VE-Cad^CREΔGrin1^ *vs* NaCl VE-Cad^CREΔGrin1^, Mann Whitney U test; see also Fig. S2B–D for comparisons across control groups).

Regarding the deep vascular plexus (Fig. 4E-G and Fig. S3), in the central region of retina, a significant increase in mesh number was observed following PAE in C57BL/6J mice (Fig. 4F; +204% for C57BL/6J, p=0.04, n=9-9, Mann Whitney U test). This effect was associated with a decrease in segment length (Fig. 4G; -13%, p=0.0244, n=9-9 for C57BL/6J, Mann Whitney U test). All these effects of PAE were not observed in VE-Cad^CREΔGrin1^ mice (Fig. 4F; +25%, p=0.3527, n=10-10 and Fig. 4G; -5%, p=0.1230, n=10-10, Mann Whitney U test). The peripheral region of the deep vascular plexus was particularly impacted by PAE in full eNMDAR mice (Fig. 4E-G). A significant increase in vascular density and mesh number were observed along with a decrease in segment length (Fig. 4E-G (blue box *vs* solid white box); 4E: +9%, p=0.0159, n=10-11; 4F: +53%, p=0.0244, n=9–9; 4G: -11%, p=0.0028, n=9–9, Mann Whitney U test). All these effects of PAE on vessel density, mesh number and segment length were abrogated in VE-Cad^CREΔGrin1^ mice (Fig. 4E-G (hatched blue box *vs* hatched white box); 4E: +4%, p=0.6607, n=10-9; 4F: +12%, p=0.3930, n=10-10; 4G: -3%, p=0.4813, n=10-10, Mann Whitney U test). In addition, the number of perforating vessels was quantified in retinal cross-sections (Fig. 4H-I). While PAE induced a significant reduction of perforating vessels in C57BL/6J mice (Fig. 4I; -24%, p=0.0093, n=8-7), this effect was not found in VE-Cad^CREΔGrin1^ mice (−13%, p=0.1490 n=5-7, Mann Whitney U test).

These data indicate that eNMDAR plays a critical role in long-term PAE-induced vascular abnormalities.

### Endothelial NMDA receptor knockout prevents PAE-induced alterations in the density of several retinal neurons

In the outer plexiform layer (OPL) to ganglion cell layer (GCL), neuronal cells are closely dependent on vascularization by retinal vascular plexuses. Therefore, we initially investigated the impact of eNMDAR knockout on the positionning of neurons located in these layers, specifically ganglion, horizontal, and amacrine cells. We examined the impact of eNMDAR knockdown on the density of these cells which were immunolabeled with RBPMS, calbindin, and calretinin antibodies, respectively. Their density was quantified in both the central and peripheral retina and results showed strain-dependent cell densities (Fig. S4). Based on these observations, the impact of eNMDAR invalidation on PAE was subsequently compared to C57BL/6J mice (Fig. 5).

**Figure 5.**
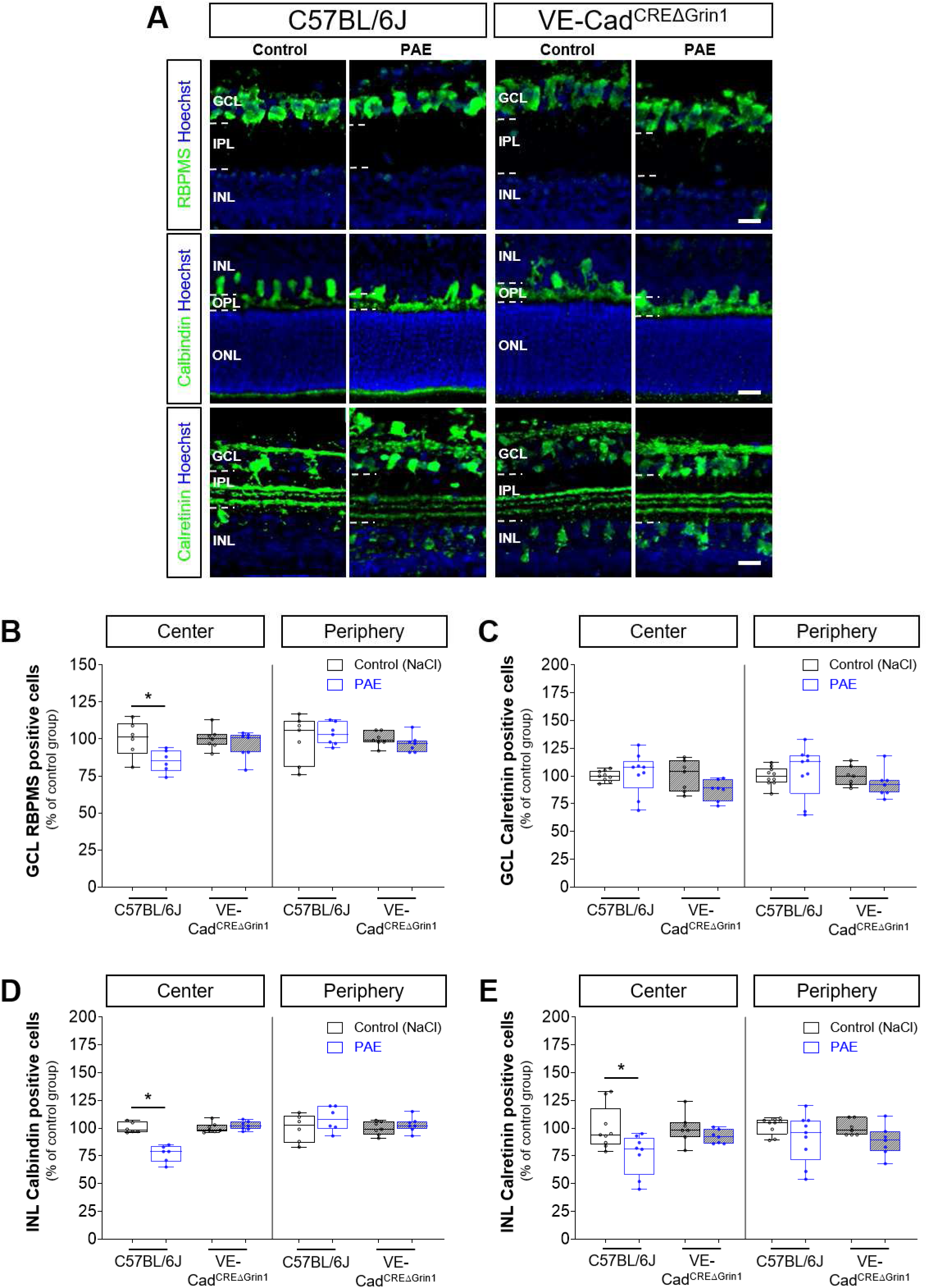
Effect of eNMDAR knockout on PAE-induced impairments on ganglion cell and interneuron densities at postnatal day 15. **(A)** Immunostaining visualizing ganglion cells (anti-RBPMS) and interneurons (anti-calbindin and anti-calretinin) in central retinal sections from C57BL/6J (solid boxes) and VE-Cad^CREΔGrin1^ (hatched boxes) mice. Nuclei are labelled with Hoechst (blue). Scale bar = 20 µm. (**B–E)** Quantification of RBPMS- (B), calretinin- (C, E) and calbindin- (D) positive cells per 100 µm in the central and peripheral retina from C57BL/6J and VE-Cad^CREΔGrin1^ mice, either prenatally exposed to alcohol (PAE, blue boxes) or not (white boxes). Data are expressed as a percentage of the mean value of the corresponding control group from the same strain. Statistical comparisons: *p ≤ 0.05 *vs* C57BL/6J control (NaCl) group, Mann–Whitney test, n = 5–7 pups per group. *GCL: Ganglion Cell Layer; INL: Inner Nuclear Layer; IPL: Inner Plexiform Layer; ONL: Outer Nuclear Layer; OPL: Outer Plexiform Layer; PAE: Prenatal Alcohol Exposure*.

In the GCL, immunolabeling with RBPMS and calretinin antibodies, which respectively label ganglion and displaced amacrine cells, revealed that PAE specifically affects ganglion cell density in central retina (Fig. 5A,B). Indeed, in GCL, PAE induced a decrease in RBPMS-positive cells (Fig. 5A) while no effect was observed for calretinin immunolabelling (Fig. 5C). Furthermore, results obtained from transgenic VE-Cad^CREΔGrin1^ mice demonstrated that eNMDAR loss prevented the PAE-induced reduction in ganglion cell density in the central retina (Fig. 5B; -15% C57BL/6J, p=0.04, n=6-6; +4% for VE-Cad^CREΔGrin1^, p=0.78, n=7-7 Mann Whitney U test).

In the inner nuclear layer (INL), a PAE-induced reduction of calbindin (Fig. 5D; -23% for C57BL/6J, p=0.0022, n=6-6, Mann Whitney U test) and calretinin-positive cells (Fig. 5E; -24% for C57BL/6J, p=0.0274, n=9-8, Mann Whitney U test) was significantly found in the central region of the retina. This effect was not observed in VE-Cad^CREΔGrin1^ mice (Fig. 5D-E (hatched blue box *vs* hatched white box); 5D: +2%, p=0.3829, n=7-7; 5E; -8%, p=0.2593, n=7-7, Mann Whitney U test).

In the outer nuclear layer (ONL), immunostaining with blue (B) and red-green (R-G) opsins and rhodopsin antibodies were performed to visualize photoreceptors (Fig. 6A). No significant effect of PAE was observed on the number of B opsin-, R-G opsin-, or rhodopsin-positive cells in either C57BL/6J or VE-Cad^CREΔGrin1^ mice (Fig. 6B–D).

**Figure 6.**
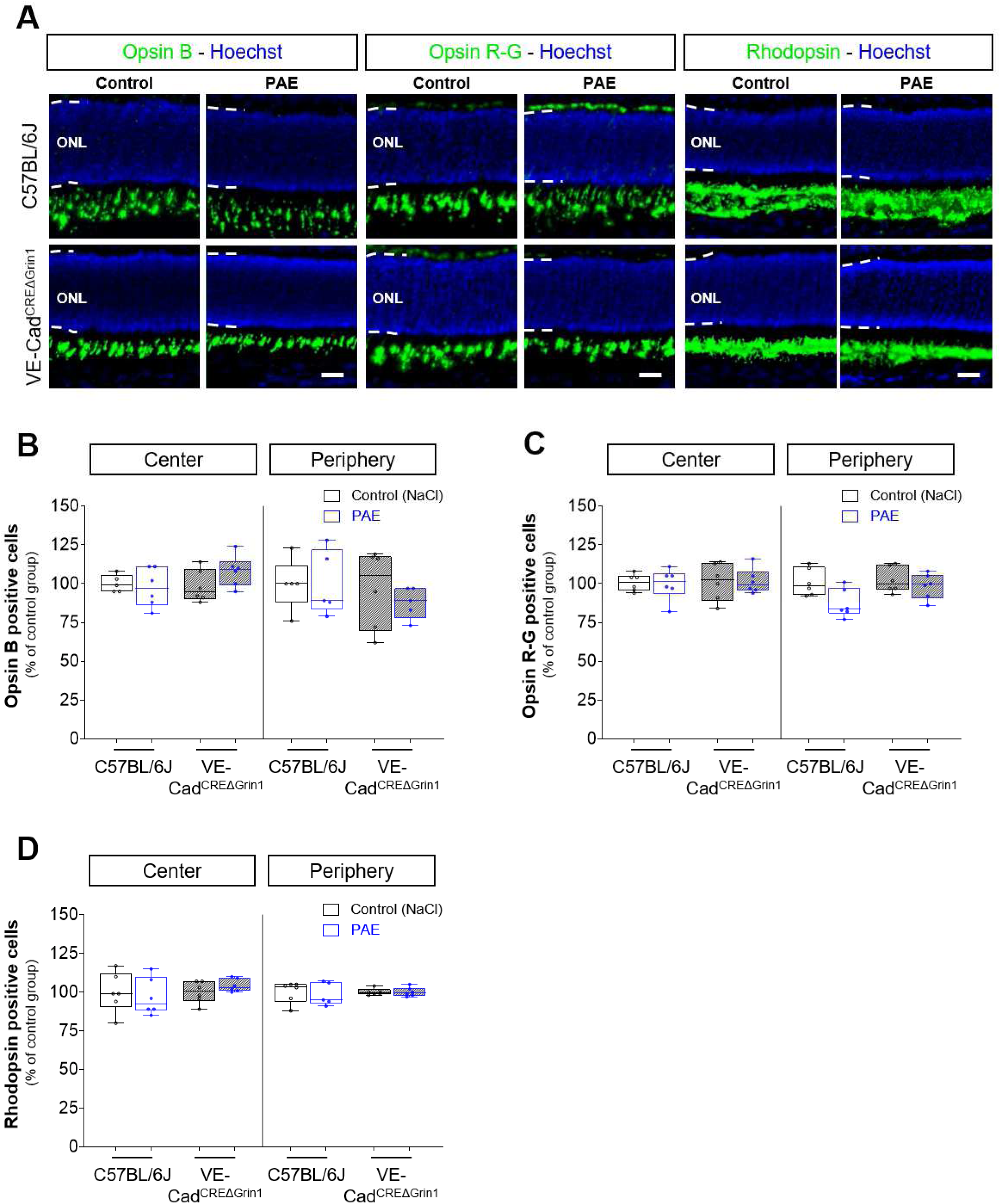
Effect of eNMDAR knockout on photoreceptors at postnatal day 15. **(A)**, Representative images showing immunohistochemistry of blue- (Opsin B), red/green- (Opsin R-G) cones, and rods (Rhodopsin) in the outer nuclear layer (ONL) on retinal sections from C57BL/6J and VE-Cad^CREΔGrin1^ mice under control (NaCl, white boxes) or PAE (blue boxes) conditions. Scale bar = 20 µm. (**B-D)** Quantification of opsin B- (B), opsin R-G- (C) and rhodopsin- (D) positive cells. Statistical comparisons: *p≤ 0.05 compared with C57BL/6J control group (NaCl). Mann-Whitney test, n=6-8 per group. *GCL: Ganglion Cell Layer; INL: Inner Nuclear Layer; IPL: Inner Plexiform Layer; ONL: Outer Nuclear Layer; OPL: Outer Plexiform Layer; PAE: Prenatal Alcohol Exposure*.

Altogether, these findings suggest that neurons positioned closer to the retinal vascular plexuses are more vulnerable to PAE, whereas photoreceptors, being not dependent on these vessels, appear less affected. Notably, the loss of eNMDAR also prevents PAE-induced alterations in these neuronal populations.

### Endothelial NMDAR knockout impairs the vascular interactions of calretinin cells at P15

Recent evidence indicates that during retinal development, calretinin-positive neurons in the INL establish close interactions with microvessels (Usui et al., 2015; Dumanoir et al., 2023). To determine whether eNMDAR loss influences these neurovascular interactions, we analyzed calretinin-positive cell associations in VE-Cad^CREΔGrin1^ mice (Fig. 7). Strikingly, VE-Cad^CREΔGrin1^ mice exhibited a significant increase in vessel-associated calretinin-positive cells compared to wild-type controls (Fig. 7B (hatched white *vs* solid white boxes); +136.1%, for VE-Cad^CREΔGrin1^ *vs* C57BL/6J and mice, n=6–8, p=0.0127). Moreover, PAE led to a substantial reduction in vessel-associated calretinin-positive cells in C57BL/6J mice (Fig. 7B (solid blue *vs* solid white box); -27.5% for PAE C57BL/6J *vs* NaCl C57BL/6J, n=7–8, p=0.0401), whereas this PAE-induced disruption was fully prevented in VE-Cad^CREΔGrin1^ mice (Fig. 7B (hatched white and blue boxes); -5.9% for PAE VE-Cad^CREΔGrin1^ *vs* NaCl VE-Cad^CREΔGrin1^, n=6–7, p=0.4452). These findings suggest that eNMDAR critically regulates the neurovascular interactions during retinal development and contributes to their vulnerability following PAE.

**Figure 7.**
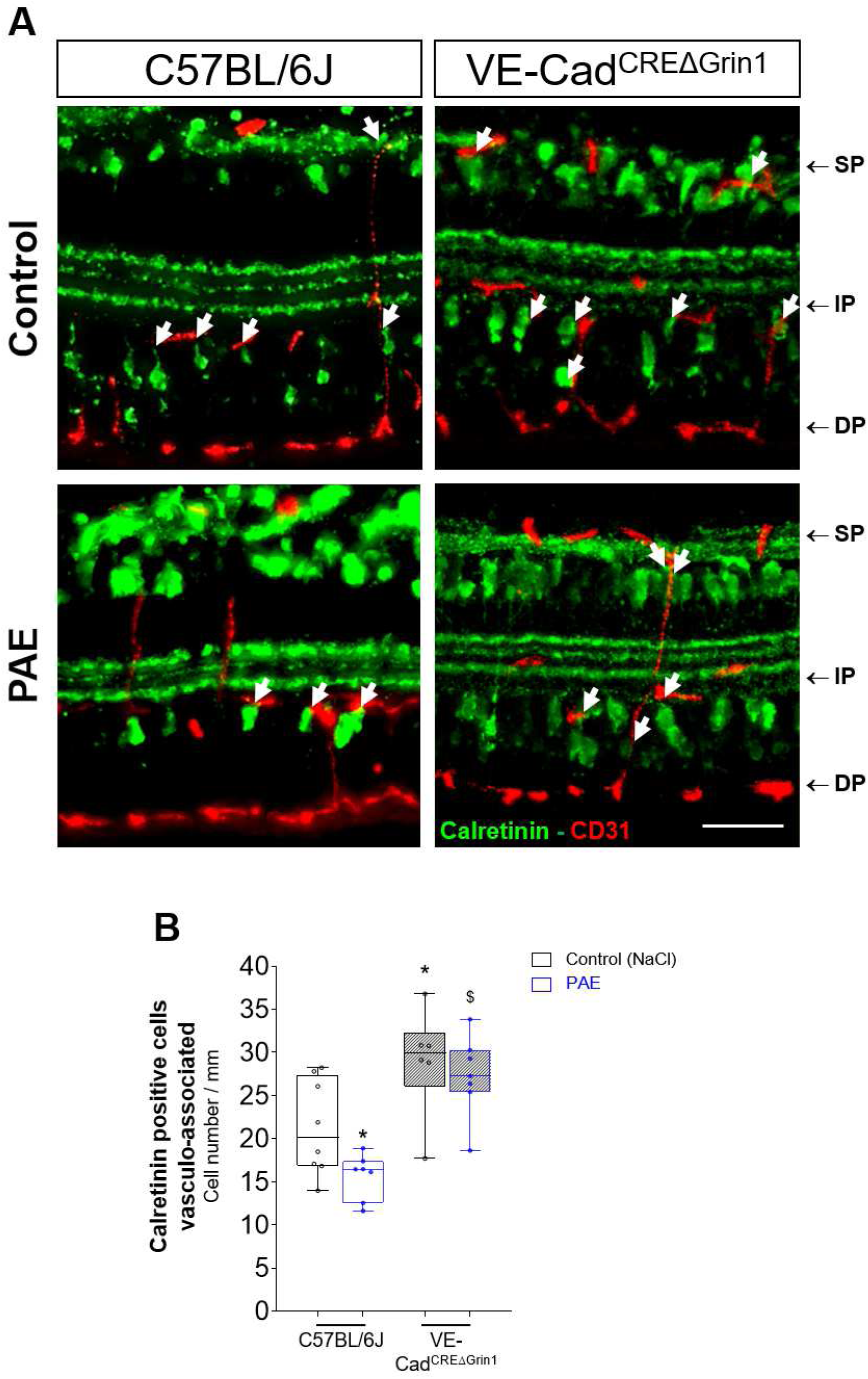
Effect of eNMDAR knockout on the density of vessel-associated calretinin-positive cells on postnatal day 15 retinas from control and PAE mice. Fluorescent immunolabeling was used to visualize calretinin-positive neurons and blood vessels (anti-CD31) in cross sections of the central retina from C57BL/6J and VE-Cad^CREΔGrin1^ mice at P15. (**A**) Representative images from control (NaCl) and PAE groups showing vessel-associated calretinin-positive cells (arrows) in the central retina of C57BL/6J and VE-Cad^CREΔGrin1^ mice. Scale bar = 20 µm. (**B**) Quantification of vessel-associated calretinin-positive cells in control (NaCl, white boxes) and PAE (blue boxes) groups for each genotype. Kruskal–Wallis test, H(3) = 16.08, p = 0.0011. Statistical comparisons: *p ≤ 0.05 *vs* C57BL/6J control (NaCl) group; $p ≤ 0.05 *vs* PAE-treated C57BL/6J group (Mann–Whitney test, n = 5–8 pups per group). *DP: Deep vascular Plexus; IP: Intermediate vascular Plexus; PAE: Prenatal Alcohol Exposure; SP: Superficial vascular Plexus*.

## Discussion

The overall results indicate that eNMDAR contributes to the development of vascular plexuses and neurogenesis in retina. Notably, eNMDAR knockout altered the vessel association of calretinin-positive interneurons. From a pathophysiological perspective, these findings suggest that eNMDAR contributes, at least partially, to PAE-induced neurovascular damage in the developing retina.

### Effect of eNMDAR knockout on retinal vascular development in mouse

Using a conditional eNMDAR knockout mouse model (VE-Cad^CREΔGrin1^) the present study demonstrated that eNMDAR invalidation impairs the development of retinal vascular plexuses. Results showed delayed progression of the superficial vascular network at P5 in VE-Cad^CREΔGrin1^ mice compared to the three other control strains in control conditions (C57BL/6J, Grin1^fl/fl^ and VE-Cad^CRE^). While the progression of the superficial network was completed by P15 in all strains, structural modifications were observed in both the superficial and deep vascular plexuses in the central and peripheral retina of VE-Cad^CREΔGrin1^ mice compared to control groups. These alterations could reflect delayed maturation of the vascular network.

Retinal vascular network development requires an initial sprouting phase to form and expand new vessels, followed by a remodeling or “pruning” phase to establish the definitive vascular mesh (Risau, 1997; Fruttiger, 2007). The vascular abnormalities observed may indicate altered development suggesting that eNMDAR could be involved in the molecular mechanisms regulating vascular sprouting and/or pruning. Supporting this hypothesis, Ferguson et al. (2016) showed that *in vitro* knockout of the GluN2D subunit in cultured endothelial cells led to reduced formation of pseudo-capillary structures, impaired endothelial cell migration, and decreased cell invasion. Similarly, Kim et al. (2022) demonstrated that in mice lacking the eNMDAR subunit 1, both vasculature and neuronal density were impaired in the neocortex, suggesting that eNMDAR deficiency result in neuronal losses. Mechanistically, interactions have been demonstrated between a major signaling pathway in angiogenesis and NMDARs (Meissirel et al., 2011). Specifically, in the developing cerebellum, VEGF, through activation of VEGF-R2, can interact and modulate NMDAR activity in cerebellar granule cells. This interaction enhances NMDAR responses, increasing intracellular calcium influx before synapse formation. Furthermore, calcium influx induced by NMDAR activation has been shown to contribute to nitric oxide production, a mediator of vasodilation and angiogenesis (Ziche and Morbidelli, 2001; Morbidelli et al., 2003; Lu et al., 2019). Taken together, these findings suggest that eNMDAR may contribute to the angiogenic process. However, the specific molecular mechanisms involved in the retina remain to be explored.

### Effect of eNMDAR invalidation on PAE-induced neurovascular damage

While the effects of *in utero* alcohol exposure on retinal development have been well established in humans (Strömland, 1987, 2004) and animal models (Clarren et al., 1990, Chmielewski et al., 1997; Tufan et al., 2007), the underlying molecular mechanisms remained unclear. Data from the present study indicate that eNMDAR knockout partially prevented PAE-induced vascular defects and abrogated alcohol’s impact on the densities of ganglion, horizontal, and amacrine cell, as well as on the number of calretinin-positive cells associated with microvessels.

The molecular mechanisms by which alcohol alters NMDAR activity have been previously investigated both *in vitro* and *in vivo* (Ren et al., 2007, 2012; Zhaoh et al., 2016). For instance, Lima-Landman (1989) demonstrated that alcohol inhibits the gating of the NMDA receptor-ion channel, reducing its open-state duration and impairing its ability to stimulate the cell. Additionally, Peoples et al. (2022) showed that the effect of ethanol on NMDAR activation kinetic depends on the specific subunit composition of the receptor. Such mechanisms contribute to the well-known effects of alcohol on human behavior, including motor incoordination and impairments in judgment and memory (Wang et al., 2024). By altering calcium channel opening, alcohol may disrupt downstream NMDAR-mediated calcium signaling pathways, thereby impacting neurovascular development. Consistent with this hypothesis, previous studies have shown that glutamate increases calcium mobilization *via* an NMDAR-dependent mechanism in cultured neonatal endothelial cells (Legros et al., 2009). In Matrigel® experiments, ethanol reduced endothelial cell tubulogenesis (Radek et al., 2008), an effect that was not observed when cells were co-incubated with MK-801, a non-competitive NMDAR antagonist. Furthermore, in organotypic slices from P2 neonates, glutamate increased calcium mobilization in cortical microvessels and the activity of endothelial MMPs, effects that were blocked by both MK-801 and ethanol (Léger et al., 2020a). Although the molecular mechanisms underlying alcohol-induced neurodevelopmental impairments are complex, multi-cellular, and far from fully understood, these findings support a role for eNMDAR in PAE-induced neurovascular damage.

### Effect of eNMDAR knockout on neurovascular interactions in mouse retina

Beyond their role in oxygen and nutrient supply, recent data suggest that vascular plexuses may also influence the interaction of developing neurons. Specifically, calretinin-positive interneurons with a migratory morphology have been shown to be closely associated with microvessels (Dumanoir et al., 2023). Data from the present study demonstrated that in VE-Cad^CREΔGrin1^ mice, density of ganglion cells and interneurons, cell types that interact closely with the superficial and deep plexuses, were altered. Notably, the number of calretinin-positive cells associated with vessels was significantly increased in VE-Cad^CREΔGrin1^ mice at P15 under control conditions. Together, these findings suggest that retinal vascular plexuses are essential for the proper development of neurons that interact with them, and that eNMDAR may play a role in regulating this vessel-dependent interaction.

During retina development, all neuronal cell types are generated from progenitor cells located at the basal pole of retina before vascular development (Amini et al., 2017). Their apical-to-basal migration occurs *via* two modes: ’somal translocation’ and ’multipolar migration’ (Amini et al., 2017). Interestingly, while vessel-associated cell migration is a well-documented process in the developing brain, particularly in the neocortex (Won et al., 2013; Tsai et al., 2016; Léger et al., 2020b), this migratory mechanism remains poorly understood in the retina. The absence of radial glia-guided migration in the retina may be explained by the fact that radial cells, such as bipolar cells and Müller glia, develop later after ganglion, amacrine, and horizontal cells (Marquardt and Gruss, 2002; Cantrup et al., 2012). Thus, in the developing superficial retinal layers, microvessels may play a supportive role in guiding the final positioning of these early-born neurons. This neurovascular interaction is thought to be critical during early postnatal development, as the architecture of the vascular network may influence neuronal migration and spatial organization. In our study, we observed that eNMDAR may contribute to modulating this interaction. Specifically, our data suggest that eNMDAR is required for the full expression of PAE-induced neurovascular alterations, indicating that eNMDARs may act as key mediators of the neurovascular crosstalk disrupted by PAE.

In conclusion, using conditional eNMDAR knockout mice and a preclinical model of FASD, this study provides the first evidence that eNMDAR is involved in the development of both the retinal vascular plexuses and several neuronal cells. These findings further suggest that glutamate, through its interaction with eNMDAR, contributes to the deleterious effects observed in FASD retinas, impacting both vascular and neuronal development. These findings provide novel insights into the molecular mechanisms by which PAE disrupts retinal vascular development, highlighting the critical role of eNMDAR. This work contributes to a better understanding of how such early vascular alterations may underlie the visual impairments commonly observed in individuals with FASD. Given the importance of the neurovascular interface in shaping retinal development and function, further studies are warranted to explore whether targeting endothelial signaling pathways, such as those involving NMDA receptors, could offer new therapeutic strategies to mitigate visual deficits associated with FASD. Longitudinal studies will also be essential to determine whether early vascular abnormalities persist into adulthood and how they correlate with long-term functional outcomes.

## Acknowledgments

This work was supported by Rouen University, Normandy University, and the French National Institute of Health and Medical Research (INSERM, UMR1245). Additional funding was provided by the Réseaux d’Intérêt Normand (RIN 3R – RIN RECHERCHE 2021-RetiGene) and the research initiative of “La Métropole Rouen Normandie” (ESR 2023). AL gratefully acknowledges the support of a fellowship from the Région Normandie.

