## supplemental figures table and legend for "Endothelial NMDA receptor involvement in retinal neurovascular damage following prenatal alcohol exposure in mouse model"

#### Supplementary figure legends

**Table S1: Statistical analyses.** Group differences were assessed using a Kruskal–Wallis test and followed Mann–Whitney tests when appropriate. The table reports the dependent variables, the factors compared, test statistics (H values and degrees of freedom), and significance levels (p values). *C: control (NaCl condition); PAE: Prenatal Alcohol Exposure.*

**Figure S1. Effect of eNMDAR knockout on the superficial and deep vascular plexus development at postnatal day 15.**

(**A**) Representative images of the superficial vascular plexuses in whole-mount retinas at P15, following CD31 immunostaining. Retinas were collected from full eNMDAR mice (i.e. C57BL/6J, Grin1$^{fl/fl}$ and VE-Cad$^{CRE}$) and eNMDAR null mice (VE-Cad$^{CRE\Delta Grin1}$ mice). Scale bars: 500 µm. Insets show high-magnification views of the superficial vascular network in the central (1) and peripheral (2) retina. Scale bars: 100 µm. (**B–D**) Quantification of the vascular density (B), mesh number (C) and segment length (D) in the central and peripheral retinas of the superficial plexus from C57BL/6J, Grin1$^{fl/fl}$, VE-Cad$^{CRE}$ (solid boxes) and VE-Cad$^{CRE\Delta Grin1}$ mice (hatched boxes) at P15. (**E–G**) Quantification of the vascular density (E), mesh number (F) and segment length (G) in the central and peripheral retinas of the deep plexus from C57BL/6J, Grin1$^{fl/fl}$, VE-Cad$^{CRE}$ (solid boxes) and VE-Cad$^{CRE\Delta Grin1}$ mice (hatched boxes) at P15. Statistical comparisons: *$p \leq 0.05$ compared with C57BL/6J control (NaCl) group, #$p \leq 0.05$ compared with Grin1$^{fl/fl}$ control (NaCl) group, \$$p \leq 0.05$ compared with VE-Cad$^{CRE}$ control (NaCl) group.

**Figure S2. Effect of eNMDAR knockout on PAE-induced alterations in the superficial vascular plexus at postnatal day 15.** (**A**) Representative images of the superficial vascular plexuses in whole-mount retinas at P15, following CD31 immunostaining. Retinas were collected from full eNMDAR mice (i.e. C57BL/6J, Grin1$^{fl/fl}$ and VE-Cad$^{CRE}$) and eNMDAR null mice (VE-Cad$^{CRE\Delta Grin1}$) mice. Scale bars: 500 µm. Insets show high-magnification images of the deep vascular network in the central (1) and periphery (2) of the retina. Scale bars: 100 µm. (**B-D**) Quantification of the vascular density (B), mesh number (C) and segment length (D) in the superficial vascular plexus at P15 in control (NaCl; white boxes) and PAE (blue boxes) retinas from C57BL/6J, Grin1$^{fl/fl}$, VE-Cad$^{CRE}$ and VE-Cad$^{CRE\Delta Grin1}$ mice in the central and peripheral retina. Results are expressed as a percent of mean of the strain matched control group (NaCl). Statistical comparisons: *p≤ 0.05 compared with strain matched control group (NaCl); Mann-Whitney test, n=7-11 per group. *PAE: Prenatal Alcohol Exposure.*

**Figure S3. Effect of eNMDAR knockout on PAE-induced alterations in the deep vascular plexus at postnatal day 15. (A)** Representative images of the deep vascular plexuses in whole-mount retinas at P15, following CD31 immunostaining. Retinas were collected from full eNMDAR mice (i.e. C57BL/6J, Grin1$^{fl/fl}$ and VE-Cad$^{CRE}$) and eNMDAR null mice (VE-Cad$^{CRE\Delta Grin1}$). Scale bars: 500 µm. Insets show high-magnification images of the deep vascular network in the central (1) and periphery (2) of the retina. Scale bars: 100 µm. (**B-D**) Quantification of the vascular density (B), mesh number (C) and segment length (D) in the deep vascular plexus at P15 in control (NaCl; white boxes) and PAE (blue boxes) retinas from C57BL/6J Grin1$^{fl/fl}$, VE-Cad$^{CRE}$ and VE-Cad$^{CRE\Delta Grin1}$ mice in the central and peripheral retina. Results are expressed as a percent of mean of the strain matched control group (NaCl). Scale bars: 200 µm. *p≤ 0.05 compared with C57BL/6J control group, #p≤ 0.05 compared with Grin1$^{fl/fl}$ control group (NaCl), $p≤ 0.05 compared with VE-Cad$^{CRE}$ control group; Mann-Whitney test, n=7-11 pups per group. *PAE: Prenatal Alcohol Exposure.*

**Figure S4. Effect of eNMDAR knockout on ganglion cells and interneurons at postnatal day 15.** Double fluorescent labeling was used to visualize neuronal cells labeled with anti-RBPMS (ganglion cells, green), anti-calbindin and anti-calretinin antibodies (interneurons, green). Nuclei were counterstained with Hoechst (blue). (**A**) Representative images showing

RBPMS-positive cells, calbindin- and calretinin-positive interneurons in central retina sections collected from full eNMDAR mice (i.e. C57BL/6J, Grin1fl/fl and VE-Cad$^{CRE}$) and eNMDAR null mice (VE-Cad$^{CRE\Delta Grin1}$). Scale bar = 20 µm. (**B-E)** Quantification of RBPMS- (B), calbindin- (D), and calretinin- (C,E) positive cells in the central and peripheral retina. Statistical comparisons: *$p \leq 0.05$ compared with C57BL/6J control (NaCl) group, #$p \leq 0.05$ compared with Grin1$^{fl/fl}$ control (NaCl) group, \$$p \leq 0.05$ compared with VECad$^{CRE}$ control (NaCl) group; Mann-Whitney test, n=6-10 pups per group. *GCL: Ganglion Cell Layer; INL: Inner Nuclear Layer; IPL: Inner Plexiform Layer; ONL: Outer Nuclear Layer; OPL: Outer Plexiform Layer; PAE: Prenatal Alcohol Exposure.*

### Table S1: Statistical analyses.

| Figure | Test | Comparisons | n | Stat Value | *p-value* |
|---|---|---|---|---|---|
| **Figure 2B** | Kruskal-Wallis test | 8 groups | 7-11 | H(7) = 28.74 | ***p=0.0002*** |
| | Mann-Whitney U test | C57BL/6J Ctrl *vs.* Grin1$^{fl/fl}$ Ctrl | 10-10 | U = 50 | *p>0.99* |
| | | C57BL/6J Ctrl *vs.* VE-Cad$^{CRE}$ Ctrl | 10-11 | U = 29 | *p=0.0720* |
| | | C57BL/6J Ctrl *vs.* VE-Cad$^{CRE\Delta Grin1}$ Ctrl | 10-9 | U = 12 | ***p=0.0057*** |
| | | Grin1$^{fl/fl}$ Ctrl *vs.* VE-Cad$^{CRE\Delta Grin1}$ Ctrl | 10-9 | U = 10 | ***p=0.0030*** |
| | | VE-Cad$^{CRE}$ Ctrl *vs.* VE-Cad$^{CRE\Delta Grin1}$ Ctrl | 11-9 | U = 18 | ***p=0.0159*** |
| | | Grin1$^{fl/fl}$ Ctrl *vs.* VE-Cad$^{CRE}$ Ctrl | 10-11 | U = 34 | *p=0.1517* |
| | | C57BL/6J Ctrl *vs.* C57BL/6J PAE | 10-10 | U = 8 | ***p=0.0007*** |
| | | Grin1$^{fl/fl}$ Ctrl *vs.* Grin1$^{fl/fl}$ PAE | 10-11 | U = 18 | ***p=0.0079*** |
| | | VE-Cad$^{CRE}$ Ctrl *vs.* VE-Cad$^{CRE}$ PAE | 11-11 | U = 28 | ***p=0.0336*** |
| | | VE-Cad$^{CRE\Delta Grin1}$ Ctrl *vs.* VE-Cad$^{CRE\Delta Grin1}$ PAE | 9-7 | U = 18 | *p=0.1738* |
| **Figure 2D** | Kruskal-Wallis test | 8 groups | 8-11 | H(7) = 42.98 | ***p<0.0001*** |
| | Mann-Whitney U test | C57BL/6J Ctrl *vs.* Grin1$^{fl/fl}$ Ctrl | 10-9 | U = 15 | ***p=0.0133*** |
| | | C57BL/6J Ctrl *vs.* VE-Cad$^{CRE}$ Ctrl | 10-10 | U = 23 | ***p=0.0433*** |
| | | C57BL/6J Ctrl *vs.* VE-Cad$^{CRE\Delta Grin1}$ Ctrl | 10-8 | U = 36 | *p=0.7618* |
| | | Grin1$^{fl/fl}$ Ctrl *vs.* VE-Cad$^{CRE\Delta Grin1}$ Ctrl | 9-8 | U = 5 | ***p=0.0016*** |
| | | VE-Cad$^{CRE}$ Ctrl *vs.* VE-Cad$^{CRE\Delta Grin1}$ Ctrl | 10-8 | U = 13 | ***p=0.0155*** |
| | | Grin1$^{fl/fl}$ Ctrl *vs.* VE-Cad$^{CRE}$ Ctrl | 9-10 | U = 42 | *p=0.8421* |
| | | C57BL/6J Ctrl *vs.* C57BL/6J PAE | 10-9 | U = 1 | ***p<0.0001*** |
| | | Grin1$^{fl/fl}$ Ctrl *vs.* Grin1$^{fl/fl}$ PAE | 9-11 | U = 23 | ***p=0.0465*** |
| | | VE-Cad$^{CRE}$ Ctrl *vs.* VE-Cad$^{CRE}$ PAE | 10-10 | U = 31 | *p=0.1655* |
| | | VE-Cad$^{CRE\Delta Grin1}$ Ctrl *vs.* VE-Cad$^{CRE\Delta Grin1}$ PAE | 8-8 | U = 21 | *p=0.2786* |
| **Figure 2E** | Kruskal-Wallis test | 8 groups | 8-11 | H(7) = 24.44 | ***p=0.0010*** |
| | Mann-Whitney U test | C57BL/6J Ctrl *vs.* Grin1$^{fl/fl}$ Ctrl | 10-9 | U = 36 | *p=0.4967* |
| | | C57BL/6J Ctrl *vs.* VE-Cad$^{CRE}$ Ctrl | 10-10 | U = 47 | *p=0.4813* |
| | | C57BL/6J Ctrl *vs.* VE-Cad$^{CRE\Delta Grin1}$ Ctrl | 10-8 | U = 35 | *p=0.6965* |
| | | Grin1$^{fl/fl}$ Ctrl *vs.* VE-Cad$^{CRE\Delta Grin1}$ Ctrl | 9-8 | U = 30 | *p=0.6058* |
| | | VE-Cad$^{CRE}$ Ctrl *vs.* VE-Cad$^{CRE\Delta Grin1}$ Ctrl | 10-8 | U = 34 | *p=0.6334* |
| | | Grin1$^{fl/fl}$ Ctrl *vs.* VE-Cad$^{CRE}$ Ctrl | 9-10 | U = 44 | *p=0.9682* |
| | | C57BL/6J Ctrl *vs.* C57BL/6J PAE | 10-9 | U = 20 | ***p=0.0435*** |
| | | Grin1$^{fl/fl}$ Ctrl *vs.* Grin1$^{fl/fl}$ PAE | 9-11 | U = 17 | ***p=0.0125*** |
| | | VE-Cad$^{CRE}$ Ctrl *vs.* VE-Cad$^{CRE}$ PAE | 10-10 | U = 12 | ***p=0.0029*** |
| | | VE-Cad$^{CRE\Delta Grin1}$ Ctrl *vs.* VE-Cad$^{CRE\Delta Grin1}$ PAE | 8-8 | U = 6 | ***p=0.0047*** |
| **Figure 2F** | Kruskal-Wallis test | 8 groups | 8-11 | H(7) = 21.04 | ***p=0.0037*** |
| | Mann-Whitney U test | C57BL/6J Ctrl *vs.* Grin1$^{fl/fl}$ Ctrl | 10-9 | U = 40 | *p=0.7197* |
| | | C57BL/6J Ctrl *vs.* VE-Cad$^{CRE}$ Ctrl | 10-10 | U = 42 | *p=0.5787* |
| | | C57BL/6J Ctrl *vs.* VE-Cad$^{CRE\Delta Grin1}$ Ctrl | 10-8 | U = 34 | *p=0.6334* |
| | | Grin1$^{fl/fl}$ Ctrl *vs.* VE-Cad$^{CRE\Delta Grin1}$ Ctrl | 9-8 | U = 26 | *p=0.3704* |
| | | VE-Cad$^{CRE}$ Ctrl *vs.* VE-Cad$^{CRE\Delta Grin1}$ Ctrl | 10-8 | U = 31 | *p=0.4598* |
| | | Grin1$^{fl/fl}$ Ctrl *vs.* VE-Cad$^{CRE}$ Ctrl | 9-10 | U = 42 | *p=0.8421* |
| | | C57BL/6J Ctrl *vs.* C57BL/6J PAE | 10-9 | U = 26 | *p=0.1333* |
| | | Grin1$^{fl/fl}$ Ctrl *vs.* Grin1$^{fl/fl}$ PAE | 9-11 | U = 16 | ***p=0.0097*** |
| | | VE-Cad$^{CRE}$ Ctrl *vs.* VE-Cad$^{CRE}$ PAE | 10-10 | U = 22 | *p=0.0355* |
| | | VE-Cad$^{CRE\Delta Grin1}$ Ctrl *vs.* VE-Cad$^{CRE\Delta Grin1}$ PAE | 8-8 | U = 8 | ***p=0.0104*** |
| **Figure 3B** | | Control *vs.* MK801 | 8-8 | t=0.7526 df=7 | *p=0.4762* |

| | | | | | |
|---|---|---|---|---|---|
| | Ratio paired *t* test | Ethanol *vs.* MK801 + Ethanol | 8-8 | t=0.0665 df=7 | *p=0.9488* |
| | | Control *vs.* Ethanol | 8-8 | t=4.858 df=7 | ***p=0.0018*** |
| | | MK801 *vs.* MK801 + Ethanol | 8-8 | t=1.849 df=7 | *p=0.1069* |
| **Figure 4B, S1B and S2B Central retina** | Kruskal-Wallis test | 8 groups | 7-11 | H(7) = 52.24 | ***p<0.0001*** |
| | Mann-Whitney U test | C57BL/6J Ctrl *vs.* Grin1$^{fl/fl}$ Ctrl | 10-7 | U = 7 | ***p=0.0046*** |
| | | C57BL/6J Ctrl *vs.* VE-Cad$^{CRE}$ Ctrl | 10-8 | U = 0 | ***p<0.0001*** |
| | | C57BL/6J Ctrl *vs.* VE-Cad$^{CRE\Delta Grin1}$ Ctrl | 10-10 | U = 46 | *p=0.7959* |
| | | Grin1$^{fl/fl}$ Ctrl *vs.* VE-Cad$^{CRE\Delta Grin1}$ Ctrl | 7-10 | U = 6 | ***p=0.0031*** |
| | | VE-Cad$^{CRE}$ Ctrl *vs.* VE-Cad$^{CRE\Delta Grin1}$ Ctrl | 8-10 | U = 0 | ***p<0.0001*** |
| | | Grin1$^{fl/fl}$ Ctrl *vs.* VE-Cad$^{CRE}$ Ctrl | 7-8 | U = 3 | ***p=0.0022*** |
| | | C57BL/6J Ctrl *vs.* C57BL/6J PAE | 10-11 | U = 26 | ***p=0.0430*** |
| | | Grin1$^{fl/fl}$ Ctrl *vs.* Grin1$^{fl/fl}$ PAE | 7-9 | U = 6 | ***p=0.0052*** |
| | | VE-Cad$^{CRE}$ Ctrl *vs.* VE-Cad$^{CRE}$ PAE | 8-7 | U = 0 | ***p=0.0003*** |
| | | VE-Cad$^{CRE\Delta Grin1}$ Ctrl *vs.* VE-Cad$^{CRE\Delta Grin1}$ PAE | 10-9 | U = 8 | ***p=0.0015*** |
| **Figure 4B, S1B and S2B Peripheral retina** | Kruskal-Wallis test | 8 groups | 7-11 | H(7) = 47.73 | ***p<0.0001*** |
| | Mann-Whitney U test | C57BL/6J Ctrl *vs.* Grin1$^{fl/fl}$ Ctrl | 10-7 | U = 3 | ***p=0.0007*** |
| | | C57BL/6J Ctrl *vs.* VE-Cad$^{CRE}$ Ctrl | 10-8 | U = 0 | ***p<0.0001*** |
| | | C57BL/6J Ctrl *vs.* VE-Cad$^{CRE\Delta Grin1}$ Ctrl | 10-10 | U = 16 | ***p=0.0089*** |
| | | Grin1$^{fl/fl}$ Ctrl *vs.* VE-Cad$^{CRE\Delta Grin1}$ Ctrl | 7-10 | U = 25 | *p=0.3638* |
| | | VE-Cad$^{CRE}$ Ctrl *vs.* VE-Cad$^{CRE\Delta Grin1}$ Ctrl | 8-10 | U = 1 | ***p<0.0001*** |
| | | Grin1$^{fl/fl}$ Ctrl *vs.* VE-Cad$^{CRE}$ Ctrl | 7-8 | U = 0 | ***p=0.0012*** |
| | | C57BL/6J Ctrl *vs.* C57BL/6J PAE | 10-11 | U = 41 | *p=0.3494* |
| | | Grin1$^{fl/fl}$ Ctrl *vs.* Grin1$^{fl/fl}$ PAE | 7-9 | U = 12 | ***p=0.0418*** |
| | | VE-Cad$^{CRE}$ Ctrl *vs.* VE-Cad$^{CRE}$ PAE | 8-7 | U = 8 | ***p=0.0205*** |
| | | VE-Cad$^{CRE\Delta Grin1}$ Ctrl *vs.* VE-Cad$^{CRE\Delta Grin1}$ PAE | 10-9 | U = 32 | *p=0.3154* |
| **Figure 4C, S1C and S2C Central retina** | Kruskal-Wallis test | 8 groups | 7-10 | H(7) = 26.87 | ***p=0.0004*** |
| | Mann-Whitney U test | C57BL/6J Ctrl *vs.* Grin1$^{fl/fl}$ Ctrl | 9-7 | U = 22 | *p=0.3510* |
| | | C57BL/6J Ctrl *vs.* VE-Cad$^{CRE}$ Ctrl | 9-7 | U = 1 | ***p=0.0003*** |
| | | C57BL/6J Ctrl *vs.* VE-Cad$^{CRE\Delta Grin1}$ Ctrl | 9-10 | U = 43 | *p=0.9048* |
| | | Grin1$^{fl/fl}$ Ctrl *vs.* VE-Cad$^{CRE\Delta Grin1}$ Ctrl | 7-10 | U = 27 | *p=0.4747* |
| | | VE-Cad$^{CRE}$ Ctrl *vs.* VE-Cad$^{CRE\Delta Grin1}$ Ctrl | 7-10 | U = 14 | ***p=0.0431*** |
| | | Grin1$^{fl/fl}$ Ctrl *vs.* VE-Cad$^{CRE}$ C | 7-7 | U = 14 | *p=0.2086* |
| | | C57BL/6J Ctrl *vs.* C57BL/6J PAE | 9-9 | U = 14 | ***p=0.0188*** |
| | | Grin1$^{fl/fl}$ Ctrl *vs.* Grin1$^{fl/fl}$ PAE | 7-8 | U = 22 | *p=0.5358* |
| | | VE-Cad$^{CRE}$ Ctrl *vs.* VE-Cad$^{CRE}$ PAE | 7-8 | U = 10 | ***p=0.0401*** |
| | | VE-Cad$^{CRE\Delta Grin1}$ Ctrl *vs.* VE-Cad$^{CRE\Delta Grin1}$ PAE | 10-10 | U = 30 | *p=0.1431* |
| **Figure 4C. S1C and S2C Peripheral retina** | Kruskal-Wallis test | 8 groups | 7-10 | H(7) = 34.50 | ***p<0.0001*** |
| | Mann-Whitney U test | C57BL/6J Ctrl *vs.* Grin1$^{fl/fl}$ Ctrl | 9-7 | U = 7 | ***p=0.0079*** |
| | | C57BL/6J Ctrl *vs.* VE-Cad$^{CRE}$ Ctrl | 9-7 | U = 0 | ***p=0.0002*** |
| | | C57BL/6J Ctrl *vs.* VE-Cad$^{CRE\Delta Grin1}$ Ctrl | 9-10 | U = 35 | *p=0.4470* |
| | | Grin1$^{fl/fl}$ Ctrl *vs.* VE-Cad$^{CRE\Delta Grin1}$ Ctrl | 7-10 | U = 14 | ***p=0.0431*** |
| | | VE-Cad$^{CRE}$ Ctrl *vs.* VE-Cad$^{CRE\Delta Grin1}$ Ctrl | 7-10 | U = 0 | ***p=0.0001*** |
| | | Grin1$^{fl/fl}$ Ctrl *vs.* VE-Cad$^{CRE}$ Ctrl | 7-7 | U = 23 | *p=0.9015* |
| | | C57BL/6J Ctrl *vs.* C57BL/6J PAE | 9-9 | U = 15 | ***p=0.0244*** |
| | | Grin1$^{fl/fl}$ Ctrl *vs.* Grin1$^{fl/fl}$ PAE | 7-8 | U = 14 | *p=0.1206* |
| | | VE-Cad$^{CRE}$ Ctrl *vs.* VE-Cad$^{CRE}$ PAE | 7-8 | U = 0 | ***p=0.0003*** |
| | | VE-Cad$^{CRE\Delta Grin1}$ Ctrl *vs.* VE-Cad$^{CRE\Delta Grin1}$ PAE | 10-10 | U = 40 | *p=0.4813* |
| | Kruskal-Wallis test | 8 groups | 9-7 | H(7) = 39.11 | ***p<0.0001*** |

| | | | | | |
|---|---|---|---|---|---|
| **Figure 4D, S1D and S2D Central retina** | Mann-Whitney U test | C57BL/6J Ctrl *vs.* Grin1$^{fl/fl}$ Ctrl | 9-7 | U = 12 | ***p=0.0418*** |
| | | C57BL/6J Ctrl *vs.* VE-Cad$^{CRE}$ Ctrl | 9-7 | U = 2 | ***p=0.0007*** |
| | | C57BL/6J Ctrl *vs.* VE-Cad$^{CRE\Delta Grin1}$ Ctrl | 9-10 | U = 37 | *P=0.5490* |
| | | Grin1$^{fl/fl}$ Ctrl *vs.* VE-Cad$^{CRE\Delta Grin1}$ Ctrl | 7-10 | U = 13 | ***p=0.03301*** |
| | | VE-Cad$^{CRE}$ Ctrl *vs.* VE-Cad$^{CRE\Delta Grin1}$ Ctrl | 7-10 | U = 5 | ***p=0.0020*** |
| | | Grin1$^{fl/fl}$ Ctrl *vs.* VE-Cad$^{CRE}$ Ctrl | 7-7 | U = 13 | *p=0.1649* |
| | | C57BL/6J Ctrl *vs.* C57BL/6J PAE | 9-9 | U = 6 | ***p=0.0012*** |
| | | Grin1$^{fl/fl}$ Ctrl *vs.* Grin1$^{fl/fl}$ PAE | 7-8 | U = 22 | *p=0.5358* |
| | | VE-Cad$^{CRE}$ Ctrl *vs.* VE-Cad$^{CRE}$ PAE | 7-8 | U = 19 | *p=0.3357* |
| | | VE-Cad$^{CRE\Delta Grin1}$ Ctrl *vs.* VE-Cad$^{CRE\Delta Grin1}$ PAE | 10-10 | U = 37 | *p=0.3527* |
| **Figure 4D, S1D and S2D Peripheral retina** | Kruskal-Wallis test | 8 groups | 9-7 | H(7) = 36.99 | ***p<0.0001*** |
| | Mann-Whitney U test | C57BL/6J Ctrl *vs.* Grin1$^{fl/fl}$ Ctrl | 9-7 | U = 0 | ***p=0.0002*** |
| | | C57BL/6J Ctrl *vs.* VE-Cad$^{CRE}$ Ctrl | 9-7 | U = 0 | ***p=0.0002*** |
| | | C57BL/6J Ctrl *vs.* VE-Cad$^{CRE\Delta Grin1}$ Ctrl | 9-10 | U = 22 | *p=0.0653* |
| | | Grin1$^{fl/fl}$ Ctrl *vs.* VE-Cad$^{CRE\Delta Grin1}$ Ctrl | 7-10 | U = 6 | ***p=0.0031*** |
| | | VE-Cad$^{CRE}$ Ctrl *vs.* VE-Cad$^{CRE\Delta Grin1}$ Ctrl | 7-10 | U = 0 | ***p=0.0001*** |
| | | Grin1$^{fl/fl}$ Ctrl *vs.* VE-Cad$^{CRE}$ Ctrl | 7-7 | U = 4 | ***p=0.0070*** |
| | | C57BL/6J Ctrl *vs.* C57BL/6J PAE | 9-9 | U = 24 | *p=0.1615* |
| | | Grin1$^{fl/fl}$ Ctrl *vs.* Grin1$^{fl/fl}$ PAE | 7-8 | U = 4 | ***p=0.0037*** |
| | | VE-Cad$^{CRE}$ Ctrl *vs.* VE-Cad$^{CRE}$ PAE | 7-8 | U = 0 | ***p=0.0003*** |
| | | VE-Cad$^{CRE\Delta Grin1}$ Ctrl *vs.* VE-Cad$^{CRE\Delta Grin1}$ PAE | 10-10 | U = 35 | *P=0.6965* |
| **Figure 4E, S1E and S3B Central retina** | Kruskal-Wallis test | 8 groups | 7-11 | H(7) = 39.36 | ***p<0.0001*** |
| | Mann-Whitney U test | C57BL/6J Ctrl *vs.* Grin1$^{fl/fl}$ Ctrl | 10-7 | U = 17 | *p=0.0878* |
| | | C57BL/6J Ctrl *vs.* VE-Cad$^{CRE}$ Ctrl | 10-8 | U = 1 | ***p<0.0001*** |
| | | C57BL/6J Ctrl *vs.* VE-Cad$^{CRE\Delta Grin1}$ Ctrl | 10-10 | U = 37 | *p=0.3527* |
| | | Grin1$^{fl/fl}$ Ctrl *vs.* VE-Cad$^{CRE\Delta Grin1}$ Ctrl | 7-10 | U = 16 | *p=0.0702* |
| | | VE-Cad$^{CRE}$ Ctrl *vs.* VE-Cad$^{CRE\Delta Grin1}$ Ctrl | 8-10 | U = 4 | ***p=0.0002*** |
| | | Grin1$^{fl/fl}$ Ctrl *vs.* VE-Cad$^{CRE}$ Ctrl | 7-8 | U = 0 | ***p=0.0003*** |
| | | C57BL/6J Ctrl *vs.* C57BL/6J PAE | 10-11 | U = 43 | *p=0.4262* |
| | | Grin1$^{fl/fl}$ Ctrl *vs.* Grin1$^{fl/fl}$ PAE | 7-8 | U = 4 | ***p=0.0037*** |
| | | VE-Cad$^{CRE}$ Ctrl *vs.* VE-Cad$^{CRE}$ PAE | 8-7 | U = 28 | *p>0.9999* |
| | | VE-Cad$^{CRE\Delta Grin1}$ Ctrl *vs.* VE-Cad$^{CRE\Delta Grin1}$ PAE | 10-9 | U = 26 | *p=0.1333* |
| **Figure 4E, S1E and S3B Peripheral retina** | Kruskal-Wallis test | 8 groups | 7-11 | H(7) = 45.62 | ***p<0.0001*** |
| | Mann-Whitney U test | C57BL/6J Ctrl *vs.* Grin1$^{fl/fl}$ Ctrl | 10-7 | U = 26 | *p=0.4173* |
| | | C57BL/6J Ctrl *vs.* VE-Cad$^{CRE}$ Ctrl | 10-8 | U = 0 | ***p<0.0001*** |
| | | C57BL/6J Ctrl *vs.* VE-Cad$^{CRE\Delta Grin1}$ Ctrl | 10-10 | U = 40 | *p=0.4813* |
| | | Grin1$^{fl/fl}$ Ctrl *vs.* VE-Cad$^{CRE\Delta Grin1}$ Ctrl | 7-10 | U = 32 | *p=0.8125* |
| | | VE-Cad$^{CRE}$ Ctrl *vs.* VE-Cad$^{CRE\Delta Grin1}$ Ctrl | 8-10 | U = 2 | ***p=0.0002*** |
| | | Grin1$^{fl/fl}$ Ctrl *vs.* VE-Cad$^{CRE}$ Ctrl | 7-8 | U = 3 | ***p=0.0022*** |
| | | C57BL/6J Ctrl *vs.* C57BL/6J PAE | 10-11 | U = 21 | ***p=0.0159*** |
| | | Grin1$^{fl/fl}$ Ctrl *vs.* Grin1$^{fl/fl}$ PAE | 7-8 | U = 10 | ***p=0.0401*** |
| | | VE-Cad$^{CRE}$ Ctrl *vs.* VE-Cad$^{CRE}$ PAE | 8-7 | U = 2 | ***p=0.0012*** |
| | | VE-Cad$^{CRE\Delta Grin1}$ Ctrl *vs.* VE-Cad$^{CRE\Delta Grin1}$ PAE | 10-9 | U = 39 | *p=0.6607* |
| **Figure 4F, S1F and S3C Central retina** | Kruskal-Wallis test | 8 groups | 7-10 | H(7) = 31.03 | ***p<0.0001*** |
| | | C57BL/6J Ctrl *vs.* Grin1$^{fl/fl}$ Ctrl | 9-7 | U = 12 | ***p=0.0418*** |
| | | C57BL/6J Ctrl *vs.* VE-Cad$^{CRE}$ Ctrl | 9-8 | U = 1 | ***p=0.0002*** |
| | | C57BL/6J Ctrl *vs.* VE-Cad$^{CRE\Delta Grin1}$ Ctrl | 9-10 | U = 37 | *p=0.5490* |
| | | Grin1$^{fl/fl}$ Ctrl *vs.* VE-Cad$^{CRE\Delta Grin1}$ Ctrl | 7-10 | U = 17 | *p=0.0878* |
| | | VE-Cad$^{CRE}$ Ctrl *vs.* VE-Cad$^{CRE\Delta Grin1}$ Ctrl | 8-10 | U = 15 | ***p=0.0266*** |
| | | Grin1$^{fl/fl}$ Ctrl *vs.* VE-Cad$^{CRE}$ Ctrl | 7-8 | U = 20 | *p=0.3969* |

| | | | | | |
|---|---|---|---|---|---|
| | Mann-Whitney U test | C57BL/6J Ctrl *vs.* C57BL/6J PAE | 9-9 | U = 17 | ***p=0.0400*** |
| | | Grin1$^{fl/fl}$ Ctrl *vs.* Grin1$^{fl/fl}$ PAE | 7-8 | U = 27 | *p=0.9551* |
| | | VE-Cad$^{CRE}$ Ctrl *vs.* VE-Cad$^{CRE}$ PAE | 8-8 | U = 10 | ***p=0.0207*** |
| | | VE-Cad$^{CRE\Delta Grin1}$ Ctrl *vs.* VE-Cad$^{CRE\Delta Grin1}$ PAE | 10-10 | U = 37 | *p=0.3927* |
| **Figure 4F, S1F and S3C Peripheral retina** | Kruskal-Wallis test | 8 groups | 7-10 | H(7) = 41.28 | ***p<0.0001*** |
| | Mann-Whitney U test | C57BL/6J Ctrl *vs.* Grin1$^{fl/fl}$ Ctrl | 9-7 | U = 16 | *p=0.1142* |
| | | C57BL/6J Ctrl *vs.* VE-Cad$^{CRE}$ Ctrl | 9-8 | U = 1 | ***p=0.0002*** |
| | | C57BL/6J Ctrl *vs.* VE-Cad$^{CRE\Delta Grin1}$ Ctrl | 9-10 | U = 40 | *p=0.7197* |
| | | Grin1$^{fl/fl}$ Ctrl *vs.* VE-Cad$^{CRE\Delta Grin1}$ Ctrl | 7-10 | U = 19 | *p=0.1331* |
| | | VE-Cad$^{CRE}$ Ctrl *vs.* VE-Cad$^{CRE\Delta Grin1}$ Ctrl | 8-10 | U = 1 | ***p<0.0001*** |
| | | Grin1$^{fl/fl}$ Ctrl *vs.* VE-Cad$^{CRE}$ Ctrl | 7-8 | U = 23 | *p=0.6126* |
| | | C57BL/6J Ctrl *vs.* C57BL/6J PAE | 9-9 | U = 15 | ***p=0.0244*** |
| | | Grin1$^{fl/fl}$ Ctrl *vs.* Grin1$^{fl/fl}$ PAE | 7-8 | U = 26 | *p=0.8665* |
| | | VE-Cad$^{CRE}$ Ctrl *vs.* VE-Cad$^{CRE}$ PAE | 8-8 | U = 8 | ***p=0.0104*** |
| | | VE-Cad$^{CRE\Delta Grin1}$ Ctrl *vs.* VE-Cad$^{CRE\Delta Grin1}$ PAE | 10-10 | U = 38 | *p=0.3930* |
| **Figure 4G, S1G and S3D Central retina** | Kruskal-Wallis test | 8 groups | 7-10 | H(7) = 37.20 | ***p<0.0001*** |
| | Mann-Whitney U test | C57BL/6J Ctrl *vs.* Grin1$^{fl/fl}$ Ctrl | 9-7 | U = 8 | ***p=0.0115*** |
| | | C57BL/6J Ctrl *vs.* VE-Cad$^{CRE}$ Ctrl | 9-8 | U = 2 | ***p=0.0003*** |
| | | C57BL/6J Ctrl *vs.* VE-Cad$^{CRE\Delta Grin1}$ Ctrl | 9-10 | U =30 | *p=0.2428* |
| | | Grin1$^{fl/fl}$ Ctrl *vs.* VE-Cad$^{CRE\Delta Grin1}$ Ctrl | 7-10 | U = 11 | ***p=0.0185*** |
| | | VE-Cad$^{CRE}$ Ctrl *vs.* VE-Cad$^{CRE\Delta Grin1}$ Ctrl | 8-10 | U = 5 | ***p=0.0009*** |
| | | Grin1$^{fl/fl}$ Ctrl *vs.* VE-Cad$^{CRE}$ Ctrl | 7-8 | U = 20 | *p=0.3969* |
| | | C57BL/6J Ctrl *vs.* C57BL/6J PAE | 9-9 | U = 15 | ***p=0.0244*** |
| | | Grin1$^{fl/fl}$ Ctrl *vs.* Grin1$^{fl/fl}$ PAE | 7-8 | U = 15 | *p=0.1520* |
| | | VE-Cad$^{CRE}$ Ctrl *vs.* VE-Cad$^{CRE}$ PAE | 8-8 | U = 7 | ***p=0.0070*** |
| | | VE-Cad$^{CRE\Delta Grin1}$ Ctrl *vs.* VE-Cad$^{CRE\Delta Grin1}$ PAE | 10-10 | U = 29 | *p=0.1230* |
| **Figure 4G, S1G and S3D Peripheral retina** | Kruskal-Wallis test | 8 groups | 7-10 | H(7) = 40.76 | ***p<0.0001*** |
| | Mann-Whitney U test | C57BL/6J Ctrl *vs.* Grin1$^{fl/fl}$ Ctrl | 9-7 | U = 15 | *p=0.0907* |
| | | C57BL/6J Ctrl *vs.* VE-Cad$^{CRE}$ Ctrl | 9-8 | U = 2 | ***p=0.0003*** |
| | | C57BL/6J Ctrl *vs.* VE-Cad$^{CRE\Delta Grin1}$ Ctrl | 9-10 | U = 21 | *p=0.2775* |
| | | Grin1$^{fl/fl}$ Ctrl *vs.* VE-Cad$^{CRE\Delta Grin1}$ Ctrl | 7-10 | U = 18 | *p=0.1088* |
| | | VE-Cad$^{CRE}$ Ctrl *vs.* VE-Cad$^{CRE\Delta Grin1}$ Ctrl | 8-10 | U = 6 | ***p=0.0014*** |
| | | Grin1$^{fl/fl}$ Ctrl *vs.* VE-Cad$^{CRE}$ Ctrl | 7-8 | U = 26 | *p=0.8665* |
| | | C57BL/6J Ctrl *vs.* C57BL/6J PAE | 9-9 | U = 8 | ***p=0.0028*** |
| | | Grin1$^{fl/fl}$ Ctrl *vs.* Grin1$^{fl/fl}$ PAE | 7-8 | U = 22 | *p=0.5358* |
| | | VE-Cad$^{CRE}$ Ctrl *vs.* VE-Cad$^{CRE}$ PAE | 8-8 | U = 7 | ***p=0.0070*** |
| | | VE-Cad$^{CRE\Delta Grin1}$ Ctrl *vs.* VE-Cad$^{CRE\Delta Grin1}$ PAE | 10-10 | U = 40 | *p=0.4813* |
| **Figure 4I** | Kruskal-Wallis test | 4 groups | 6-10 | H(3) = 41.22 | ***p=0.0204*** |
| | Mann-Whitney U test | C57BL/6J Ctrl *vs.* VE-Cad$^{CRE\Delta Grin1}$ Ctrl | 8-5 | U = 10 | *p=0.1709* |
| | | C57BL/6J PAE *vs.* VE-Cad$^{CRE\Delta Grin1}$ PAE | 7-7 | U = 16 | *p=0.3176* |
| | | C57BL/6J Ctrl *vs.* C57BL/6J PAE | 8-7 | U = 6 | ***p=0.0093*** |
| | | VE-Cad$^{CRE\Delta Grin1}$ Ctrl *vs.* VE-Cad$^{CRE\Delta Grin1}$ PAE | 5-7 | U = 8 | *p=0.1490* |
| **Figure 5B Central retina** | Kruskal-Wallis test | 4 groups | 6-7 | H(3) = 8.048 | ***p=0.0450*** |
| | Mann-Whitney U test | C57BL/6J Ctrl *vs.* C57BL/6J PAE | 6-6 | U = 5 | ***p=0.0411*** |
| | | VE-Cad$^{CRE\Delta Grin1}$ Ctrl *vs.* VE-Cad$^{CRE\Delta Grin1}$ PAE | 7-7 | U = 22 | *p=0.7815* |
| **Figure 5B peripheral retina** | Kruskal-Wallis test | 4 groups | 7-7 | H(3) = 8.048 | *p=0.0783* |

| | | | | | |
|---|---|---|---|---|---|
| **Figure 5C Central retina** | Kruskal-Wallis test | 4 groups | 7-9 | H(3) = 3.655 | *p=0.3012* |
| **Figure 5C peripheral retina** | Kruskal-Wallis test | 4 groups | 7-9 | H(3) = 3.657 | *p=0.3010* |
| **Figure 5D Central retina** | Kruskal-Wallis test | 4 groups | 6-7 | H(3) = 17.64 | ***p=0.0005*** |
| | Mann-Whitney U test | C57BL/6J Ctrl *vs.* C57BL/6J PAE | 6-6 | U = 0 | ***p=0.0022*** |
| | | VE-Cad$^{CRE\Delta Grin1}$ Ctrl *vs.* VE-Cad$^{CRE\Delta Grin1}$ PAE | 7-7 | U = 17 | *p=0.3829* |
| **Figure 5D peripheral retina** | Kruskal-Wallis test | 4 groups | 6-7 | H(3) = 9.27 | ***p=0.0259*** |
| | Mann-Whitney U test | C57BL/6J Ctrl *vs.* C57BL/6J PAE | 6-6 | U = 11 | *p=0.3095* |
| | | VE-Cad$^{CRE\Delta Grin1}$ Ctrl *vs.* VE-Cad$^{CRE\Delta Grin1}$ PAE | 7-7 | U = 19 | *p=0.5146* |
| **Figure 5E Central retina** | Kruskal-Wallis test | 4 groups | 7-9 | H(3) = 16.02 | ***p=0.0011*** |
| | Mann-Whitney U test | C57BL/6J Ctrl *vs.* C57BL/6J PAE | 9-8 | U = 13.5 | ***p=0.0274*** |
| | | VE-Cad$^{CRE\Delta Grin1}$ Ctrl *vs.* VE-Cad$^{CRE\Delta Grin1}$ PAE | 7-7 | U = 14.5 | *p=0.2593* |
| **Figure 5E peripheral retina** | Kruskal-Wallis test | 4 groups | 7-9 | H(3) = 12.90 | ***p=0.0049*** |
| | Mann-Whitney U test | C57BL/6J Ctrl *vs.* C57BL/6J PAE | 9-9 | U = 29 | *p=0.3401* |
| | | VE-Cad$^{CRE\Delta Grin1}$ Ctrl *vs.* VE-Cad$^{CRE\Delta Grin1}$ PAE | 7-7 | U = 10 | *p=0.0728* |
| **Figure S4B Central retina** | Kruskal-Wallis test | 4 groups | 5-7 | H(3) = 17.06 | ***p=0.0007*** |
| | Mann-Whitney U test | C57BL/6J Ctrl *vs.* Grin1$^{fl/fl}$ Ctrl | 6-5 | U = 0 | ***p=0.0043*** |
| | | C57BL/6J Ctrl *vs.* VE-Cad$^{CRE}$ Ctrl | 6-6 | U = 18 | p>0.9999 |
| | | C57BL/6J Ctrl *vs.* VE-Cad$^{CRE\Delta Grin1}$ Ctrl | 6-7 | U = 6 | ***p=0.0350*** |
| | | Grin1$^{fl/fl}$ Ctrl *vs.* VE-Cad$^{CRE\Delta Grin1}$ Ctrl | 5-7 | U = 1 | ***p=0.0051*** |
| | | VE-Cad$^{CRE}$ *vs.* VE-Cad$^{CRE\Delta Grin1}$ Ctrl | 6-7 | U = 0 | ***p=0.0012*** |
| | | Grin1$^{fl/fl}$ Ctrl *vs.* VE-Cad$^{CRE}$ Ctrl | 5-6 | U = 0 | ***p=0.0043*** |
| **Figure S4B peripheral retina** | Kruskal-Wallis test | 4 groups | 5-7 | H(3) = 15.65 | ***p=0.0013*** |
| | Mann-Whitney U test | C57BL/6J Ctrl *vs.* Grin1$^{fl/fl}$ Ctrl | 7-5 | U = 17 | p>0.9999 |
| | | C57BL/6J Ctrl *vs.* VE-Cad$^{CRE}$ C | 7-7 | U = 0 | ***p=0.0006*** |
| | | C57BL/6J Ctrl *vs.* VE-Cad$^{CRE\Delta Grin1}$ Ctrl | 7-7 | U = 16 | p=0.3176 |
| | | Grin1$^{fl/fl}$ Ctrl *vs.* VE-Cad$^{CRE\Delta Grin1}$ Ctrl | 5-7 | U = 11 | p=0.3434 |
| | | VE-Cad$^{CRE}$ Ctrl *vs.* VE-Cad$^{CRE\Delta Grin1}$ Ctrl | 7-7 | U = 0 | ***p=0.0006*** |
| | | Grin1$^{fl/fl}$ Ctrl *vs.* VE-Cad$^{CRE}$ Ctrl | 5-7 | U = 0 | ***p=0.0025*** |
| **Figure S4C Central retina** | Kruskal-Wallis test | 4 groups | 6-9 | H(3) = 17.32 | ***p=0.0006*** |
| | Mann-Whitney U test | C57BL/6J Ctrl *vs.* Grin1$^{fl/fl}$ Ctrl | 9-7 | U = 3 | ***p=0.0012*** |
| | | C57BL/6J Ctrl *vs.* VE-Cad$^{CRE}$ Ctrl | 9-6 | U = 6 | ***p=0.0120*** |
| | | C57BL/6J Ctrl *vs.* VE-Cad$^{CRE\Delta Grin1}$ Ctrl | 9-7 | U = 20 | *p=0.2523* |
| | | Grin1$^{fl/fl}$ Ctrl *vs.* VE-Cad$^{CRE\Delta Grin1}$ Ctrl | 7-7 | U = 2 | ***p=0.0023*** |
| | | VE-Cad$^{CRE}$ Ctrl *vs.* VE-Cad$^{CRE\Delta Grin1}$ Ctrl | 6-7 | U = 13 | *p=0.2949* |
| | | Grin1$^{fl/fl}$ Ctrl *vs.* VE-Cad$^{CRE}$ Ctrl | 7-6 | U = 0 | ***p=0.0012*** |
| **Figure S4C peripheral retina** | Kruskal-Wallis test | 4 groups | 6-9 | H(3) = 17.91 | ***p=0.0005*** |
| | Mann-Whitney U test | C57BL/6J Ctrl *vs.* Grin1$^{fl/fl}$ Ctrl | 9-6 | U = 5 | ***p=0.0076*** |
| | | C57BL/6J Ctrl *vs.* VE-Cad$^{CRE}$ Ctrl | 9-6 | U = 5 | ***p=0.0076*** |
| | | C57BL/6J Ctrl *vs.* VE-Cad$^{CRE\Delta Grin1}$ Ctrl | 9-7 | U = 16 | p=0.1142 |

| | | | | | |
|---|---|---|---|---|---|
| | | Grin1$^{fl/fl}$ Ctrl *vs.* VE-Cad$^{CRE\Delta Grin1}$ Ctrl | 6-7 | U = 0 | ***p=0.0012*** |
| | | VE-Cad$^{CRE}$ Ctrl *vs.* VE-Cad$^{CRE\Delta Grin1}$ Ctrl | 6-7 | U = 11 | *p=0.1807* |
| | | Grin1$^{fl/fl}$ Ctrl *vs.* VE-Cad$^{CRE}$ Ctrl | 6-6 | U = 0 | ***p=0.0022*** |
| **Figure S4D Central retina** | Kruskal-Wallis test | 4 groups | 6-7 | H(3) = 5.57 | *p=0.1344* |
| **Figure S4D peripheral retina** | Kruskal-Wallis test | 4 groups | 6-7 | H(3) = 11.05 | ***p=0.015*** |
| | Mann-Whitney U test | C57BL/6J Ctrl *vs.* Grin1$^{fl/fl}$ Ctrl | 6-6 | U = 16 | p=0.8182 |
| | | C57BL/6J Ctrl *vs.* VE-Cad$^{CRE}$ Ctrl | 6-6 | U = 13 | *p=0.4848* |
| | | C57BL/6J Ctrl *vs.* VE-Cad$^{CRE\Delta Grin1}$ Ctrl | 6-7 | U = 6 | ***p=0.0350*** |
| | | Grin1$^{fl/fl}$ Ctrl C *vs.* VE-Cad$^{CRE\Delta Grin1}$ Ctrl | 6-7 | U = 5 | ***p=0.0221*** |
| | | VE-Cad$^{CRE}$ Ctrl *vs.* VE-Cad$^{CRE\Delta Grin1}$ Ctrl | 6-7 | U = 0 | ***p=0.0012*** |
| | | Grin1$^{fl/fl}$ Ctrl *vs.* VE-Cad$^{CRE}$ Ctrl | 6-6 | U = 11 | *p=0.3095* |
| **Figure S4E Central retina** | Kruskal-Wallis test | 4 groups | 6-7 | H(3) = 8.575 | ***p=0.035*** |
| | Mann-Whitney U test | C57BL/6J Ctrl *vs.* Grin1$^{fl/fl}$ Ctrl | 9-7 | U = 20 | *p=0.2523* |
| | | C57BL/6J Ctrl *vs.* VE-Cad$^{CRE}$ Ctrl | 9-6 | U = 21 | p=0.5287 |
| | | C57BL/6J Ctrl *vs.* VE-Cad$^{CRE\Delta Grin1}$ Ctrl | 9-7 | U = 15 | *p=0.0907* |
| | | Grin1$^{fl/fl}$ Ctrl *vs.* VE-Cad$^{CRE\Delta Grin1}$ Ctrl | 7-7 | U = 4 | ***p=0.0070*** |
| | | VE-Cad$^{CRE}$ Ctrl *vs.* VE-Cad$^{CRE\Delta Grin1}$ Ctrl | 6-7 | U = 12 | *p=0.2343* |
| | | Grin1$^{fl/fl}$ Ctrl *vs.* VE-Cad$^{CRE}$ Ctrl | 7-6 | U = 8 | *p=0.0734* |
| **Figure S4E peripheral retina** | Kruskal-Wallis test | 4 groups | 6-7 | H(3) = 22.27 | ***p<0.0001*** |
| | Mann-Whitney U test | C57BL/6J Ctrl *vs.* Grin1$^{fl/fl}$ Ctrl | 9-6 | U = 12 | p=0.0879 |
| | | C57BL/6J Ctrl *vs.* VE-Cad$^{CRE}$ Ctrl | 9-6 | U = 0 | ***p=0.0004*** |
| | | C57BL/6J Ctrl *vs.* VE-Cad$^{CRE\Delta Grin1}$ Ctrl | 9-7 | U = 0 | ***p=0.0002*** |
| | | Grin1$^{fl/fl}$ Ctrl *vs.* VE-Cad$^{CRE\Delta Grin1}$ Ctrl | 6-7 | U = 0 | ***p=0.0012*** |
| | | VE-Cad$^{CRE}$ Ctrl *vs.* VE-Cad$^{CRE\Delta Grin1}$ Ctrl | 6-7 | U = 5 | ***p=0.0221*** |
| | | Grin1$^{fl/fl}$ Ctrl *vs.* VE-Cad$^{CRE}$ Ctrl | 6-6 | U = 0 | ***p=0.0022*** |
| **Figure 6B Central retina** | Kruskal-Wallis test | 4 groups | 5-7 | H(3) = 1.747 | *p=0.6265* |
| **Figure 6B peripheral retina** | Kruskal-Wallis test | 4 groups | 5-6 | H(3) = 0.9455 | *p=0.8144* |
| **Figure 6C Central retina** | Kruskal-Wallis test | 4 groups | 6-7 | H(3) = 1.970 | *p=0.5798* |
| **Figure 6C peripheral retina** | Kruskal-Wallis test | 4 groups | 6-6 | H(3) = 3.988 | *p=0.2628* |
| **Figure 6D Central retina** | Kruskal-Wallis test | 4 groups | 6-7 | H(3) = 13.65 | ***p=0.034*** |
| | Mann-Whitney U test | C57BL/6J Ctrl *vs.* C57BL/6J PAE | 6-6 | U = 14 | *p=0.5649* |
| | | VE-Cad$^{CRE\Delta Grin1}$ Ctrl *vs.* VE-Cad$^{CRE\Delta Grin1}$ PAE | 6-7 | U = 18.5 | *p=0.7576* |
| **Figure 6D peripheral retina** | Kruskal-Wallis test | 4 groups | 5-6 | H(3) = 16.05 | ***p=0.0011*** |
| | Mann-Whitney U test | C57BL/6J Ctrl Ctrl *vs.* C57BL/6J PAE | 6-5 | U = 15 | *p>0.99* |
| | | VE-Cad$^{CRE\Delta Grin1}$ Ctrl *vs.* VE-Cad$^{CRE\Delta Grin1}$ PAE | 6-6 | U = 17 | *p=0.8918* |
| **Figure 7B** | Kruskal-Wallis test | 4 groups | 6-8 | H(3) = 16.08 | ***p=0.0011*** |
| | | C57BL/6J Ctrl *vs.* VE-Cad$^{CRE\Delta Grin1}$ Ctrl | 8-6 | U = 5 | ***p=0.0127*** |
| | | VE-Cad$^{CRE\Delta Grin1}$ PAE *vs.* VE-Cad$^{CRE\Delta Grin1}$ PAE | 7-7 | U = 1 | ***p=0.0012*** |

| | | | | | |
|---|---|---|---|---|---|
| | Mann-Whitney U test | C57BL/6J Ctrl *vs.* C57BL/6J PAE | 8-7 | U = 10 | ***p=0.0401*** |
| | | VE-Cad$^{CRE\Delta Grin1}$ Ctrl *vs.* VE-Cad$^{CRE\Delta Grin1}$ PAE | 6-7 | U = 15 | *p=0.4452* |

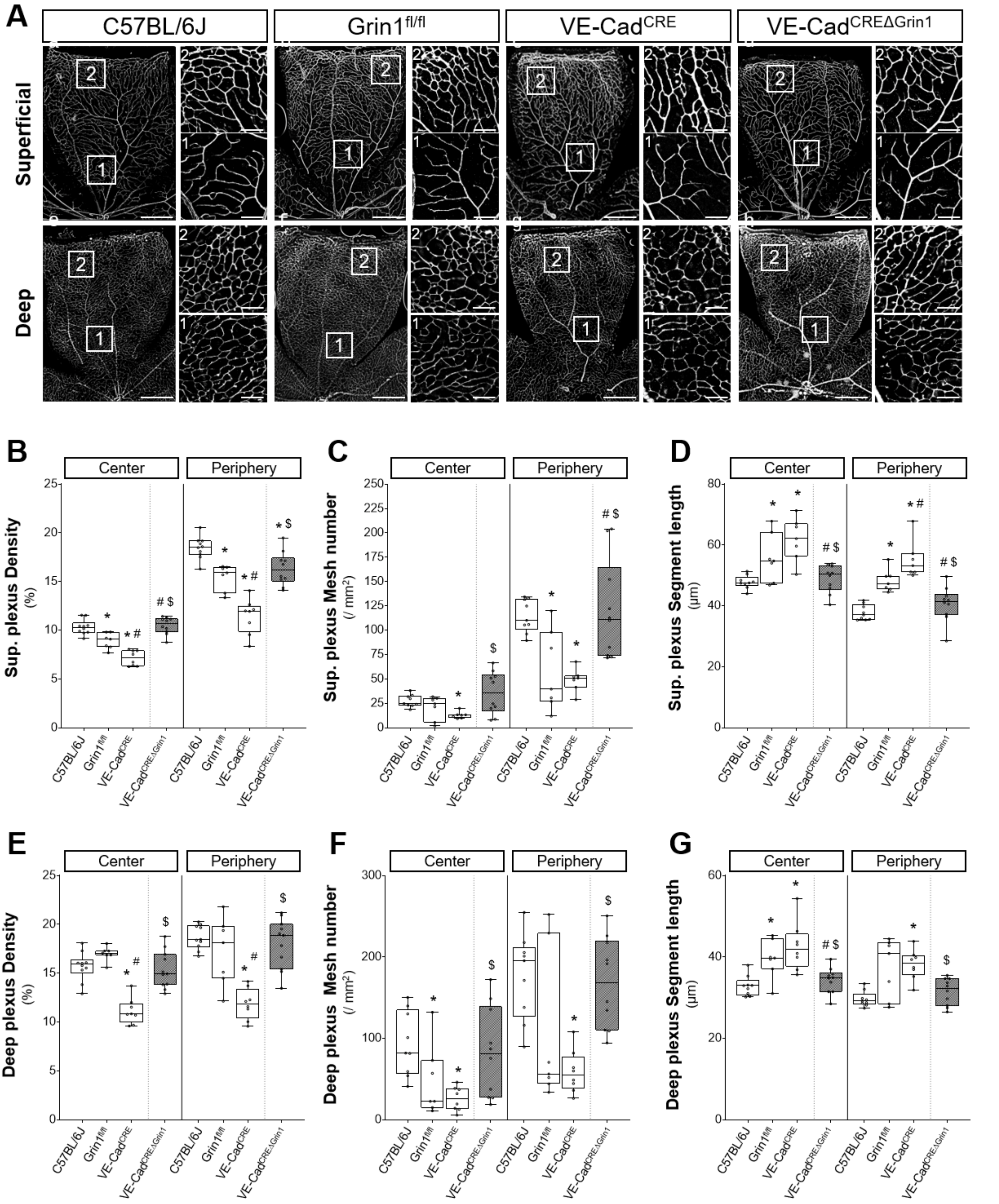


Figure S1, Leroy et al.

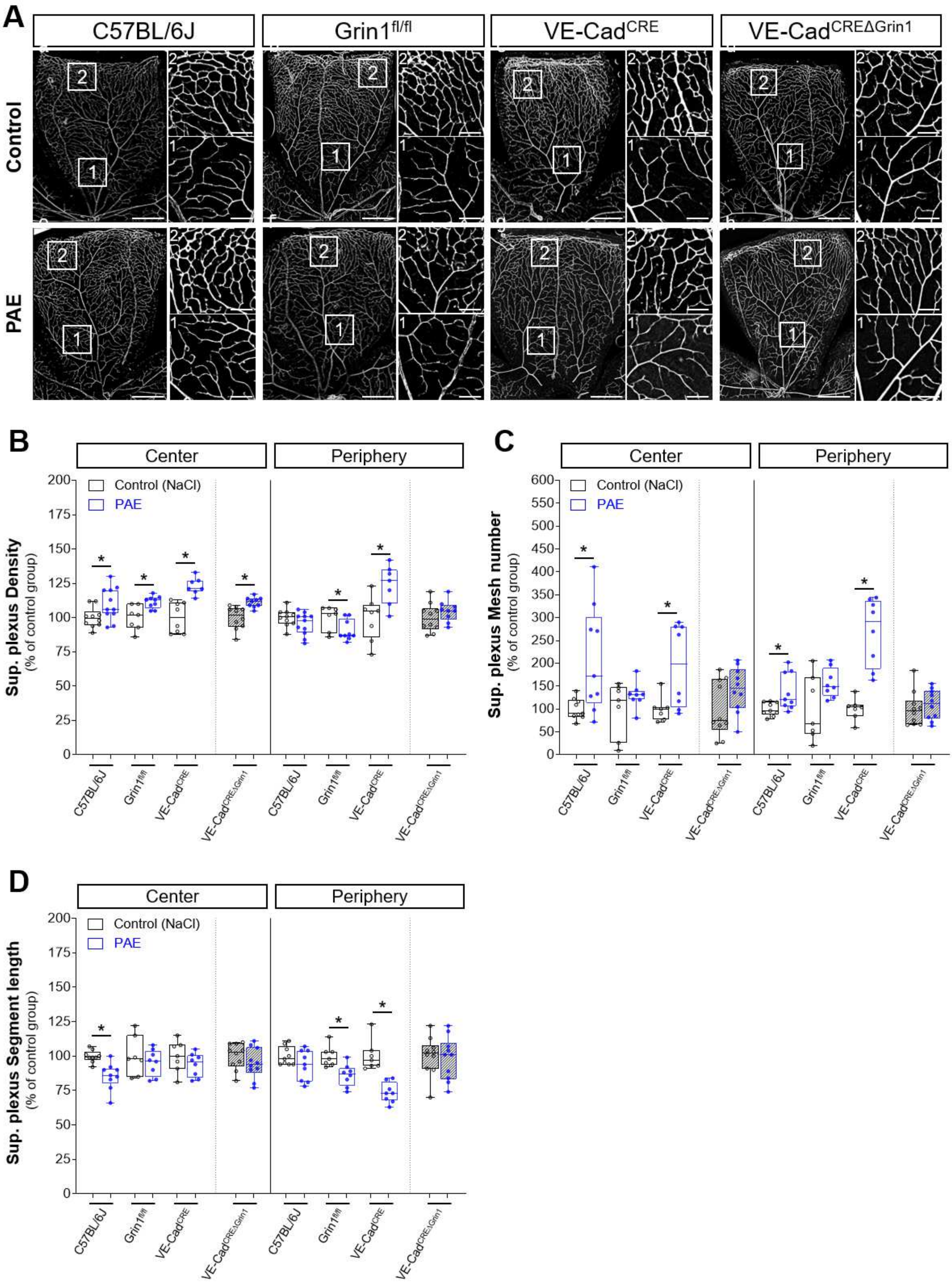


Figure S2, Leroy et al.

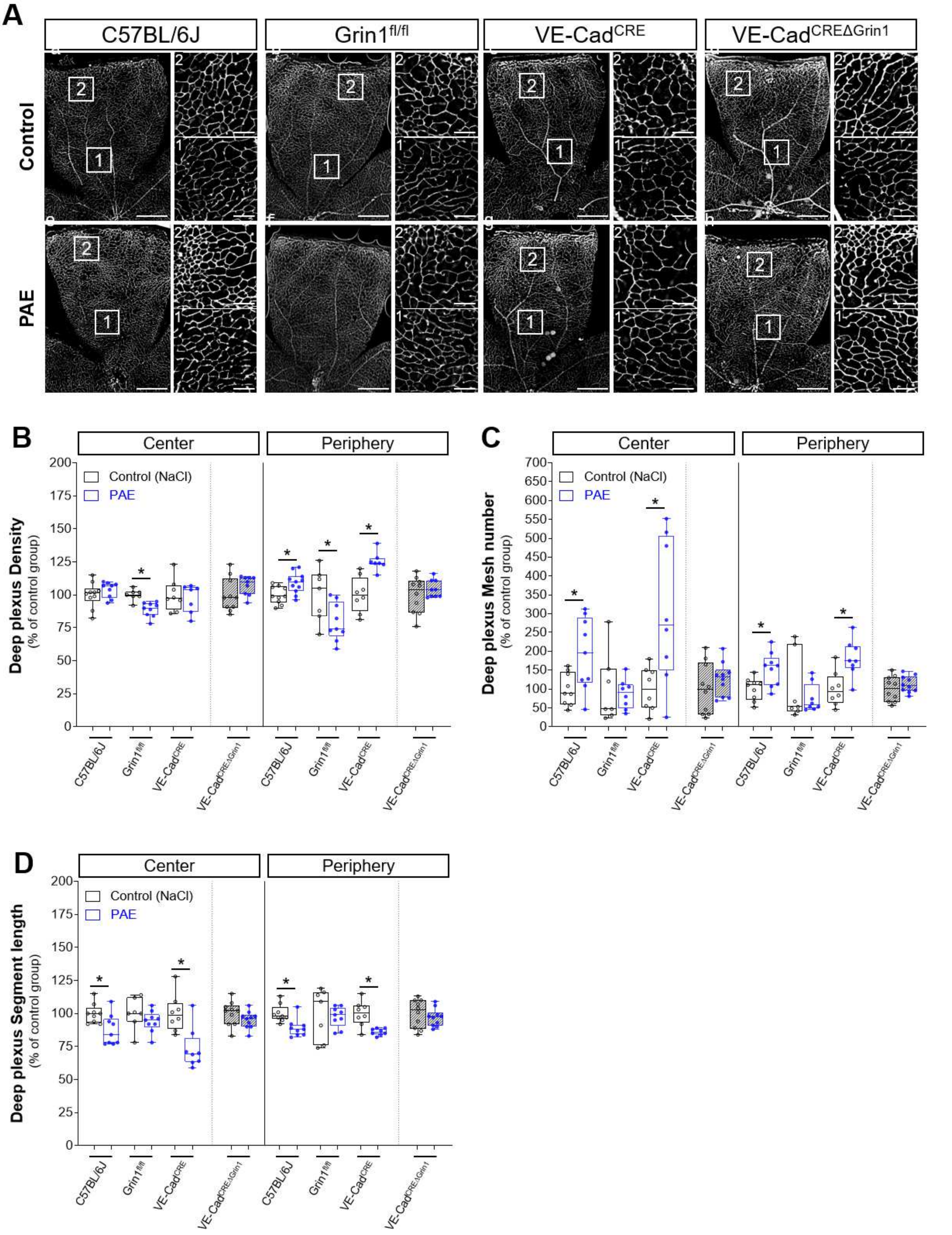


**Figure S3, Leroy et al.**

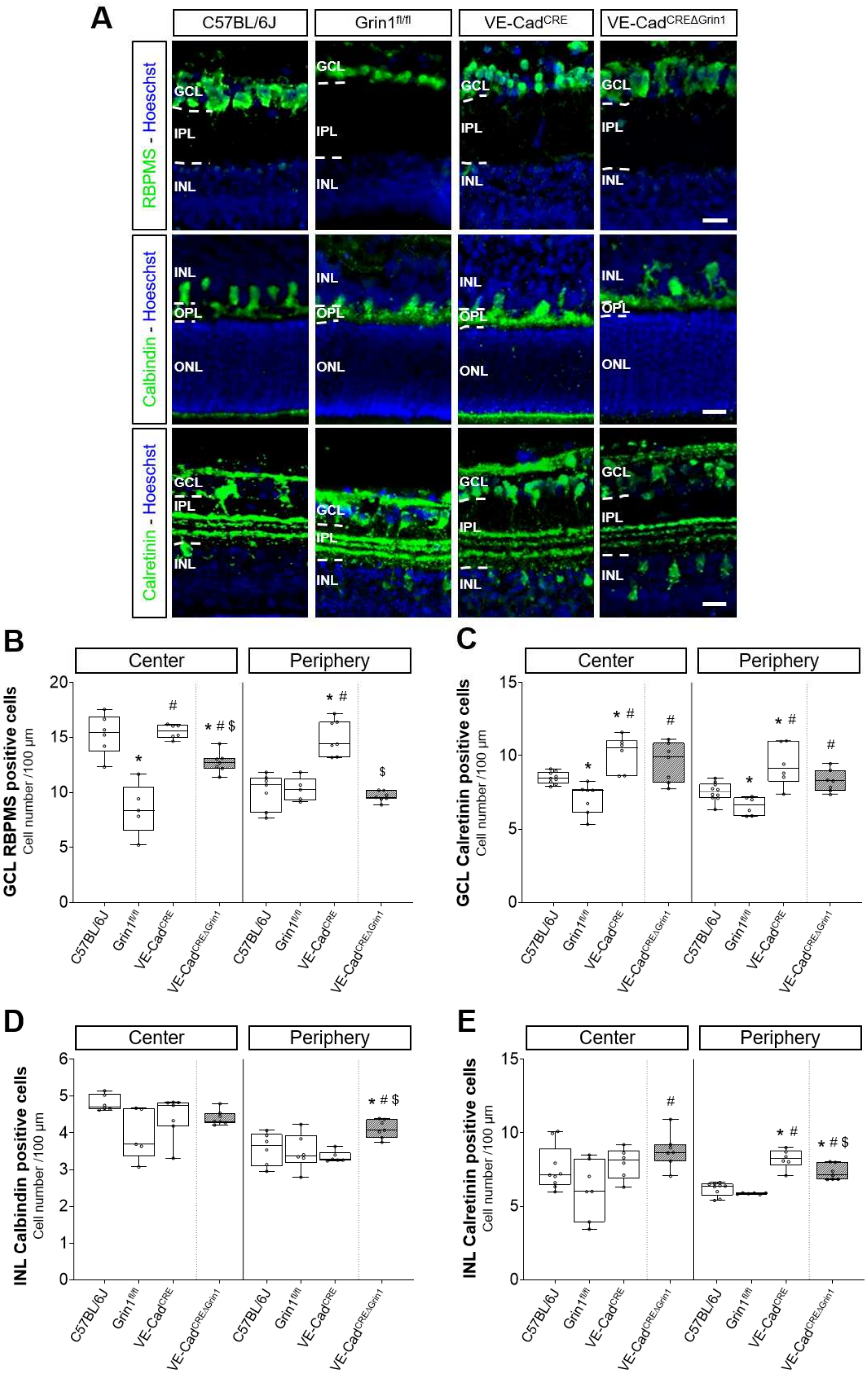


Figure S4, Leroy et al.
